# Characterization of factors that underlie transcriptional silencing in *C. elegans* oocytes

**DOI:** 10.1101/2022.08.28.505591

**Authors:** Mezmur D. Belew, Emilie Chien, W. Matthew Michael

## Abstract

While it has been appreciated for decades that prophase-arrested oocytes are transcriptionally silenced on a global level, the molecular pathways that promote silencing have remained elusive. Previous work in *C. elegans* has shown that both topoisomerase II (TOP-2) and condensin II collaborate with the H3K9me heterochromatin pathway to silence gene expression in the germline during L1 starvation, and that the PIE-1 protein silences the genome in the P-lineage of early embryos. Here, we show that all three of these silencing systems, TOP-2/condensin II, H3K9me, and PIE-1, are required for transcriptional repression in oocytes. We find that H3K9me3 marks increase dramatically on chromatin during silencing, and that silencing is under cell cycle control. We also find that PIE-1 localizes to the nucleolus just prior to silencing, and that nucleolar dissolution during silencing is dependent on TOP-2/condensin II. Our data identify both the molecular components and the trigger for genome silencing in oocytes and establish a link between PIE-1 nucleolar residency and its ability to repress transcription.

## Introduction

Cells can control gene expression at many levels, including individual loci, groups of genes that perform a common function, or at the level of the entire genome. Whole-genome control of transcription occurs in many different contexts. For example, cells undergoing quiescence silence their genomes (Swygert and Tsukiyama, 2019; de Morree and Rando, 2023), as do all proliferating cells as they prepare for mitosis (Palozola et al., 2019; Gonzalez et al., 2021; Ito and Zaret, 2022). In addition, during germline development, many animals silence the genome in germline progenitor cells until they have been specified as germline (Nakamura and Seydoux, 2008; Wang and Seydoux, 2013). Furthermore, a common feature of gametogenesis is a genome silencing event as gametes complete meiotic prophase (Schultz et al., 2018; Tora and Vincent, 2021). The genome can also be globally activated, and examples of this are also found in early development, where many organisms undergo a zygotic genome activation event as embryos transition from maternal to zygotic control of embryogenesis (Kobayashi and Tachibana, 2021).

While the concept of whole-genome activation and silencing has been appreciated for a long time, it is only recently that the molecular mechanisms in play have begun to be elucidated. One particularly useful model system for the study of whole-genome control of transcription has been the roundworm *C. elegans*. Remarkably, during germline development in the worm, the genome is silenced and then reactivated on at least four distinct occasions. During early embryogenesis, in one- and two-cell embryos, transcription is globally repressed via binding of the OMA-1 and OMA-2 proteins to the basal transcription factor TAF-4, as this sequesters TAF-4 in the cytoplasm (Guven-Ozkan et al., 2008). At the four-cell stage somatic genomes are activated. However, in germline precursor cells, the so-called P-lineage, transcription remains globally silenced (Mello et al., 1996; Seydoux et al., 1996). Silencing in the P-lineage is controlled by PIE-1, a zinc-finger containing protein whose mechanism of action during silencing is not yet fully understood. Upon germline specification, PIE-1 is degraded and germline genome activation occurs in the Z2 and Z3 primordial germ cells (PGCs; Schaner et al., 2003). PGCs remain transcriptionally active through hatching of the embryo into an L1 larva. Recent work from our group has shown that if embryos hatch in an environment lacking nutrients, then the PGC genome is silenced once again (Belew et al., 2021). Starvation-induced genome silencing is triggered by the energy sensing kinase AMPK and requires the TOP-2/condensin II chromatin compaction pathway as well as components of the H3K9me/heterochromatin pathway (Belew et al., 2021). In this system, TOP-2/condensin II promotes H3K9me2 and -me3 deposition on germline chromatin, and thus we named this pathway the Global Chromatin Compaction (GCC) to reflect the linear organization of the system components. We have also previously shown that when starved L1s are fed, then the germline genome is reactivated. This requires the induction of DNA double-strand breaks that serve to promote chromatin decompaction (Butuči et al., 2015; Wong et al., 2018). Germline chromatin remains transcriptionally active through the remainder of development and, in hermaphrodites, through oogenesis until the genome is silenced once again at the end of meiotic prophase (Schisa et al., 2001; Walker et al., 2007). Thus, in the nematode germline, there are four genome silencing events - in early embryos by OMA-1/2, in the P-lineage by PIE-1, in starved L1s by the GCC pathway, and in oocytes via an unknown mechanism.

The global repression of transcription during late oogenesis is not unique to *C. elegans*. In *Drosophila melanogaster*, oocytes are developmentally arrested at prophase I and repress transcription between the fifth and eighth stages of oogenesis, just before their re-entry into meiosis (Navarro-Costa et al., 2016). Similarly, in mice, primary follicles that contain prophase I arrested oocytes have been shown to repress transcription just before the resumption of meiosis (Moore et al., 1974; Schultz et al., 2018; Tora and Vincent, 2021). This repression persists throughout fertilization until minor ZGA occurs at first cleavage (Moore and Lintern-Moore, 1974; Abe et al., 2018). The conservation of this theme across organisms of different complexity suggests that global transcriptional repression in oocytes is an important feature of the oocyte-to-embryo transition. Despite this conservation, however, the molecular pathway(s) responsible for repression are still unknown.

In this study, we address the problem of how oocytes in *C. elegans* shut down transcription as a function of completing meiotic prophase. The nematode is an ideal system to address this important question, as the events leading up to oocyte maturation and fertilization are well described. During germline development in hermaphrodites, animals first produce sperm and then switch to making oocytes (Schedl, 1997). Oocytes are produced in an assembly-line like process, where cells exit pachytene and then progress through the remainder of meiotic prophase within the proximal portion of the tube-shaped gonad (Greenstein, 2005). Within the proximal gonad, oocytes can be clearly identified by their position relative to the spermatheca — the oocyte closest is named −1, and the next most proximal oocyte −2, et cetera. Previous work has shown that −4 and −3 oocytes are transcriptionally active, and then the genome is silenced at the −2 position (Walker et al., 2007; Figure 1A). Other work has shown that, just prior to genome silencing, the condensin II complex is recruited to chromatin, where it helps to compact chromatin during the formation of bivalents, a unique chromosome structure that enables the subsequent meiotic divisions (Chan et al., 2004; see Figure 1A). By the −2 position, bivalents have formed and the genome is silenced. At the −1 position the oocyte receives a signal from sperm to initiate maturation, and the cells then enter meiotic M-phase (Miller et al., 2001; Figure 1A). Here, we show that genome silencing in oocytes is organized by cyclin-dependent kinase 1 (CDK-1 in *C. elegans*) and requires the known silencers TOP-2/condensin II, the H3K9me/heterochromatin pathway, and PIE-1. Loss of any one of these components results in aberrant RNA polymerase II (RNAPII) activity at the −2 position. Interestingly, we also report that in oocytes distal to the −2 position, PIE-1 is mainly localized in the nucleolus, and that at the −2 position the nucleolus dissolves in a TOP-2/condensin II dependent manner. Our data identify the molecular components for the oocyte genome silencing system and suggest a model where the nucleolar residency of PIE-1 prevents it from blocking RNAPII activity until the nucleolus dissolves at the −2 position.

**Figure 1:**
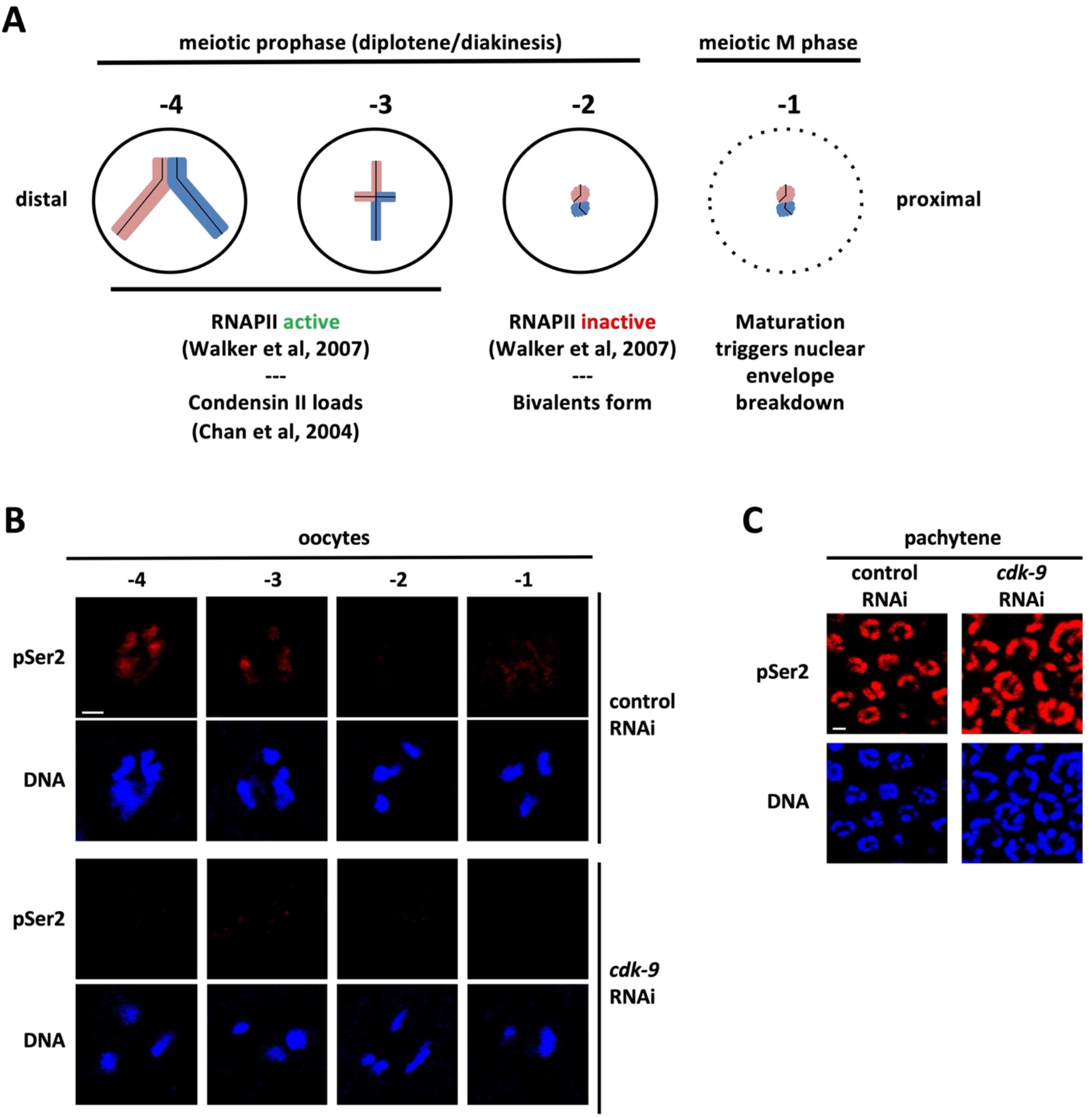
Timing and CDK-9 dependence of the phosphorylation of RNAPIISer2 in *C. elegans* proximal oocytes. A. Schematic summarizing transcriptional activity and chromatin compaction in the four most proximal oocytes. See Introduction for details. B. N2s were treated with either control or *cdk-9* RNAi. Dissected gonads from these animals were fixed and stained for DNA (blue) and RNAPIIpSer2 (red). Depletion of CDK-9 results in the loss of RNAPIIpSer2 signal in proximal oocytes. Scale bar represents a length of 2 µm. C. Pachytene nuclei from the same animals in (B) were fixed and stained for DNA (blue) and RNAPIIpSer2 (red). Unlike proximal oocytes *cdk-9* RNAi does not alter RNAPIIpSer2 signal in pachytene nuclei. Scale bar represents a length of 2 µm.

## Results

### RNAPIIpSer2 is dependent on CDK-9 in proximal oocytes

The goal of this study was to analyze genome silencing during oogenesis in *C. elegans*. To monitor transcription, we used an antibody that recognizes the active and elongating form of RNAPII. This reagent is a rabbit polyclonal antibody termed ab5095, purchased from Abcam (Waltham, MA), and it detects a phospho-epitope on the second serine within the carboxy-terminal repeat domain (CTD) of RNAPII (RNAPIIpSer2; Palancade and Bensaude, 2003). In previous work we had validated ab5095 for use in *C. elegans* via the demonstration that reactivity depends on the presence of both RNAPII and phosphate, and that ab5095 accurately labels transcriptionally active nuclei in worm embryos and in the PGCs of L1 larvae (Belew et al., 2021). For the current study we used it to examine the transcriptional status of oocytes obtained from gonads dissected from adult hermaphrodites. Prior work that examined proximal oocytes has shown that RNAPIIpSer2 signals decreased starting with the fourth most proximal oocyte (termed −4) and became undetectable through the two most proximal oocytes (−2 and −1) (Walker et al., 2007). We, therefore, focused our analysis on the four most proximal oocytes and found that RNAPIIpSer2 signal was detected invariably on the chromatin of oocytes at the −4 position (Figure 1B, control RNAi panel). However, as we examined the three most proximal oocytes, we observed several different patterns for the RNAPIIpSer2 signal: (1) signal exclusively on chromatin, (2) no signal at all, (3) signal present both on and off chromatin, and (4) a nucleoplasmic RNAPIIpSer2 signal that mostly excludes chromatin (Figures 1B and S1A&B). It thus appears that while in some oocytes RNAPIIpSer2 signal completely disappeared, in others, it was merely removed from the chromatin and was present in the nucleoplasm. This form of nucleoplasmic signal has also been observed in human cells as they approach mitosis and is explained by RNAPII coming off the chromatin while maintaining CTD phosphorylations (Hintermair et al., 2016).

In mammals, the kinase that phosphorylates serine 2 within the RNAPII CTD is P-TEFb, which is composed of the CDK9 kinase and cyclins T1 or T2 (Fujinaga et al., 2023). To determine if the corresponding kinase in *C. elegans*, CDK-9, plays a similar role in proximal oocytes, we used RNAi to deplete the protein and we then stained for RNAPIIpSer2. As shown in Figures 1B and S1A&B, exposure to *cdk-9* RNAi caused a reduction in all three forms of RNAPIIpSer2 signal (on chromatin, off chromatin, and both on and off chromatin). This shows that the off-chromatin signals are due to CDK-9 phosphorylated RNAPII that had been displaced from chromatin but not yet dephosphorylated. For the remainder of this study, we only considered nuclei with RNAPIIpSer2 on chromatin as being transcriptionally active, and the off-chromatin signals were ignored. Interestingly, we also noticed that in more distal regions of the gonad, for example the pachytene region, RNAPIIpSer2 signal was unchanged by *cdk-9* RNAi (Figure 1C). This is consistent with previous work showing that CDK-12 is the major RNAPIIpSer2 kinase in the mitotic and pachytene regions of the gonad (Bowman et al., 2013). It thus appears that there is a handoff, from CDK-12 to CDK-9, for generating RNAPIIpSer2 during oogenesis in *C. elegans*.

### The TOP-2/condensin II axis controls genome silencing during meiotic prophase

Our data, together with previous findings (Walker et al., 2007), show that genome silencing initiates at the −3 position and is largely complete by the −2 position. As detailed above, there are four genome silencing systems that have been identified thus far in *C. elegans*: the TOP-2/condensin II axis (in starved L1 PGCs; Belew et al., 2021), the H3K9me pathway (in starved L1 PGCs; Belew et al., 2021); the OMA-1/2 proteins (in one- and two-cell embryos; Guven-Ozkan et al., 2008), and the PIE-1 protein (in the P2, P3, and P4 embryonic blastomeres; Mello et al., 1996; Seydoux et al., 1996). Which, if any, of these systems might also be used for genome silencing in oocytes? We know that OMA-1/2 are dispensable for oocyte silencing, as previous work had directly examined this possibility (Guven-Ozkan et al., 2008). Thus, we turned our attention to the remaining three systems and focused initially on TOP-2/condensin II. To do so we examined RNAPIIpSer2 in oocytes from animals exposed to either *top-2, capg-2* (which encodes a protein specific for the condensin II complex), or double *top-2/capg-2* RNAi. All three of these treatments impacted the RNAPIIpSer2 pattern in a similar manner: signal now persisted in −2 oocytes, indicating that genome silencing had failed (Figure 2A&B). To quantify these data, we measured RNAPIIpSer2 signal intensity present on chromatin within individual oocytes. We normalized these values against signal intensities present in the pachytene region, as these values do not change upon depletion of TOP-2 and/or CAPG-2 (data not shown). This was done to account for potential differences in antibody penetration across the different sample sets (see Methods for a more detailed explanation). As shown in Figure 2B, in the control samples, there was a significant difference in RNAPIIpSer2 signal intensity between the −3 and −2 positions, reflecting the genome silencing that occurs in −2 oocytes under normal conditions. Also shown in Figure 2B is the difference in RNAPIIpSer2 signal intensity between −2 oocytes from control samples, relative to the samples where TOP-2 and/or CAPG-2 had been depleted. Loss of TOP-2/CAPG-2 caused a significant increase in RNAPIIpSer2 signal at the −2 position. Based on these data, we conclude that loss of TOP-2 and/or CAPG-2 impacts genome silencing in −2 oocytes. Since genome silencing normally occurs in late prophase, one possibility is that loss of TOP-2/CAPG-2 somehow delays progression through diakinesis, and thus the persistence of RNAPII transcription in −2 oocytes of *top-2/capg-2* depleted samples is an indirect consequence of defects in prophase timing. This is not the case, however, as when we stained for the prophase marker phospho-serine 10 of histone H3 (H3pS10; Hendzel et al., 1997) we saw no difference in the timing of H3pS10 deposition (Figure S2). We note that H3pS10 signal intensity was seemingly increased after *top-2/capg-2* RNAi, however in the absence of an independent source of H3pS10 signal that can be used to normalize signal intensity across different sample sets it is impossible to know if this is a physiological effect or an artifact of differential antibody penetration. Nonetheless, it is clear that depletion of TOP-2/CAPG-2 does not disrupt prophase timing.

**Figure 2:**
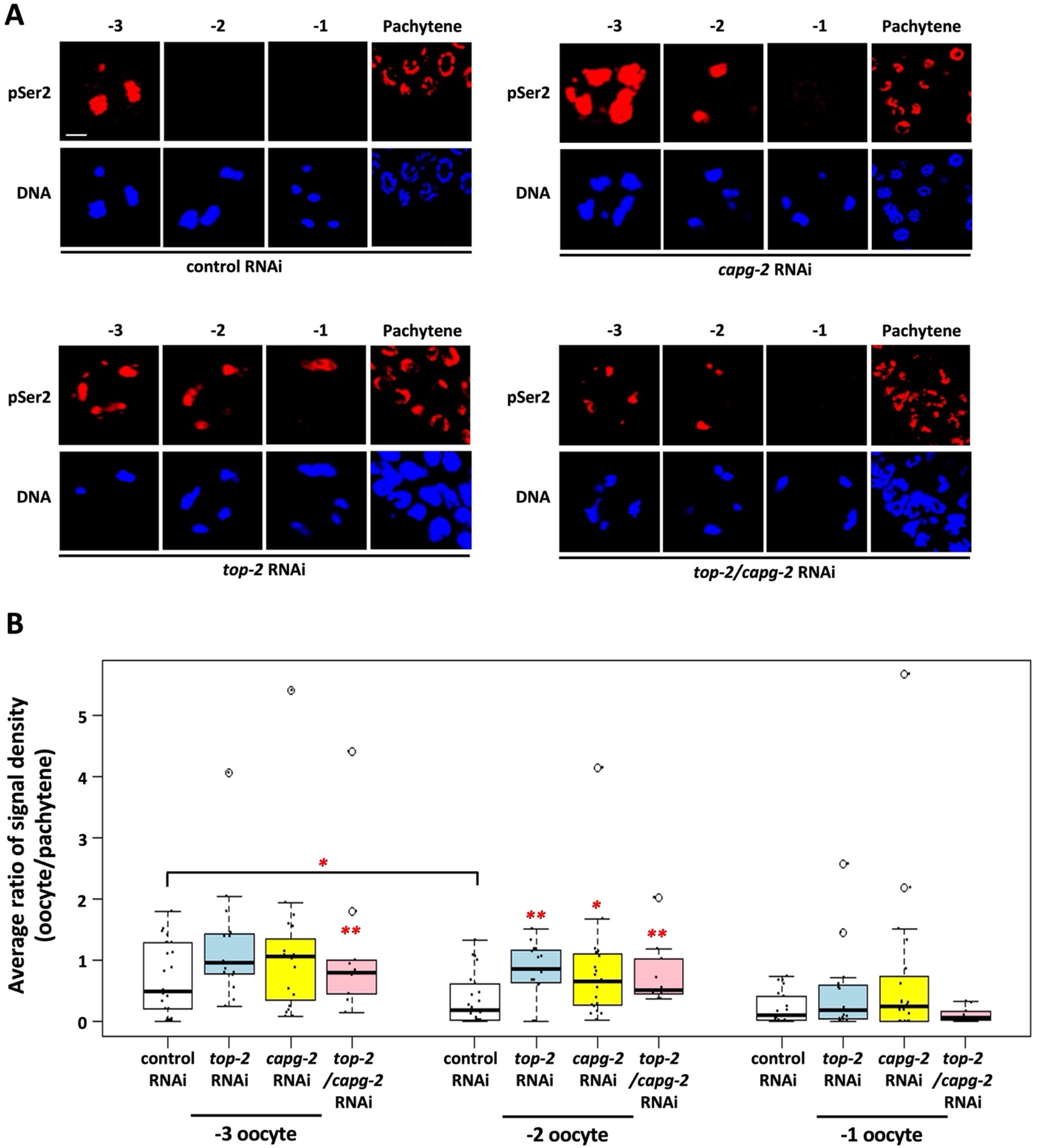
Aberrant RNAPIIpSer2 signal is observed in proximal oocytes when TOP-2 and condensin II are depleted. A. N2 animals were treated with control, *top-2*, *capg-2*, or *top-2/capg-2* double RNAi. Gonads were dissected from these animals and were fixed and stained for DNA (blue) and RNAPIIpSer2 (red). Each panel series (−3, −2, −1, and pachytene) represent nuclei from the same gonad. Individual depletion of TOP-2, CAPG-2, and their co-depletion results in persistent RNAPIIpSer2 signal on the chromatin of −3 and −2 oocytes. Scale bar represents a length of 2 µm. B. Quantification of data presented in (A). RNAPIIpSer2 signal was measured using ImageJ and values were obtained for both oocyte and pachytene nuclei. The pachytene values were then used to normalize oocyte values (please see Methods), and the average ratio of normalized raw integrated density is plotted on the y-axis. Between 10 and 22 gonads were analyzed per condition and data were collected across multiple independently performed replicates. For control RNAi 22 gonads across 5 replicates were analyzed, for *top-2* RNAi 15 gonads were analyzed across 2 replicates, for *capg-2* RNAi it was 20 gonads across 2 replicates and for *top-2/capg-2* RNAi it was 10 gonads across a single replicate. Significance was measured using student’s t-test or Wilcoxon Rank Sum test. **p<0.01; *p<0.05. In control animals, the ratio of signal density decreases significantly between −3 and −2 oocyte positions. Loss of TOP-2 or CAPG-2 results in persistent RNAPIIpSer2 signal in −2 oocytes. Co-depletion of TOP-2 and CAPG-2 produces a stronger phenotype, with persistent RNAPIIpSer2 signal in −3 and −2 oocytes.

### CDK-1 acts upstream of TOP-2/condensin II to trigger genome silencing

Our data identify a role for TOP-2 and condensin II in the silencing of transcription as oocytes prepare for maturation and the meiotic divisions. Silencing initiates at the −3 position and is complete by −2, and this is also when condensin II is recruited to oocyte chromatin (Chan et al., 2004). Previous work has also shown that condensin II activity is dependent on phosphorylation by CDK1 (Abe et al., 2011), and thus it stands to reason that CDK-1 may also play a role in genome silencing, through regulation of condensin II. To test this hypothesis, we used RNAi to reduce CDK-1 activity in proximal oocytes and we assessed the impact on transcriptional activity. As shown in Figures 3A&B, *cdk-1* RNAi triggered a significant increase in RNAPIIpSer2 signal intensity in −3, −2, and −1 oocytes, showing that CDK-1 does indeed control genome silencing. To pursue this further, we next asked the opposite question – how does elevating CDK-1 activity impact genome silencing? For this we targeted the CDK-1 inhibitory kinase WEE-1.3 for depletion, as previous work has shown that WEE-1.3 negatively regulates CDK-1 in nematode oocytes and, further, that CDK-1 is the sole target of WEE-1.3 (Burrows et al., 2006). Treatment with *wee-1.3* RNAi resulted in a decrease in RNAPIIpSer2 signal intensity across all oocytes spanning the −6 to −2 positions, showing that genome silencing was happening much earlier than in the control condition (Figures 4A&B). These data posed an important question: is the precocious silencing observed after *wee-1.3* RNAi also dependent on TOP-2/condensin II? To address this, we co-depleted WEE-1.3 and CAPG-2, and we found that transcriptional activity was now restored to proximal oocytes (Figures 4A&B). Thus, co-depletion with CAPG-2 reverses the effects of WEE-1.3 depletion alone, and this shows that condensin II is required for the precocious silencing observed when CDK-1 levels are increased via depletion of WEE-1.3. Based on these data, we conclude that CDK-1 acts upstream of TOP-2/condensin II, in a positive manner, to promote genome silencing. Furthermore, our data suggest that CDK-1 can activate condensin II for genome compaction and silencing at an activity state below that needed to trigger nuclear envelope breakdown (NEB) and entry into meiotic M-phase. This, in turn, suggests that CDK-1 activity rises gradually in the proximal gonad, as opposed to an abrupt activation in −1 oocytes. This would be consistent with previous work in HeLa cells as well as frog egg extracts showing that CDK-1-cyclin B activity rises gradually during interphase and prophase (Gavet and Pines, 2010; Maryu and Yang, 2022). To gain additional evidence for a gradual acquisition of the M-phase state in proximal oocytes we stained them with MPM-2, an antibody that recognizes mitotic phosphoproteins (Davis et al., 1983). As shown in Figure S3, MPM-2 antigens are present at low levels in −5 oocytes and they gradually accumulate as we move proximally in the gonad, such that −4 and −3 oocytes show an intermediate level while −2 and −1 oocytes show a high level of MPM-2 reactivity. These data are consistent with the idea that CDK-1 activity gradually increases, and that levels of CDK-1 that are proficient to trigger genome compaction and silencing in −3/−2 oocytes are not yet sufficient for NEB and entry into meiotic M-phase.

**Figure 3:**
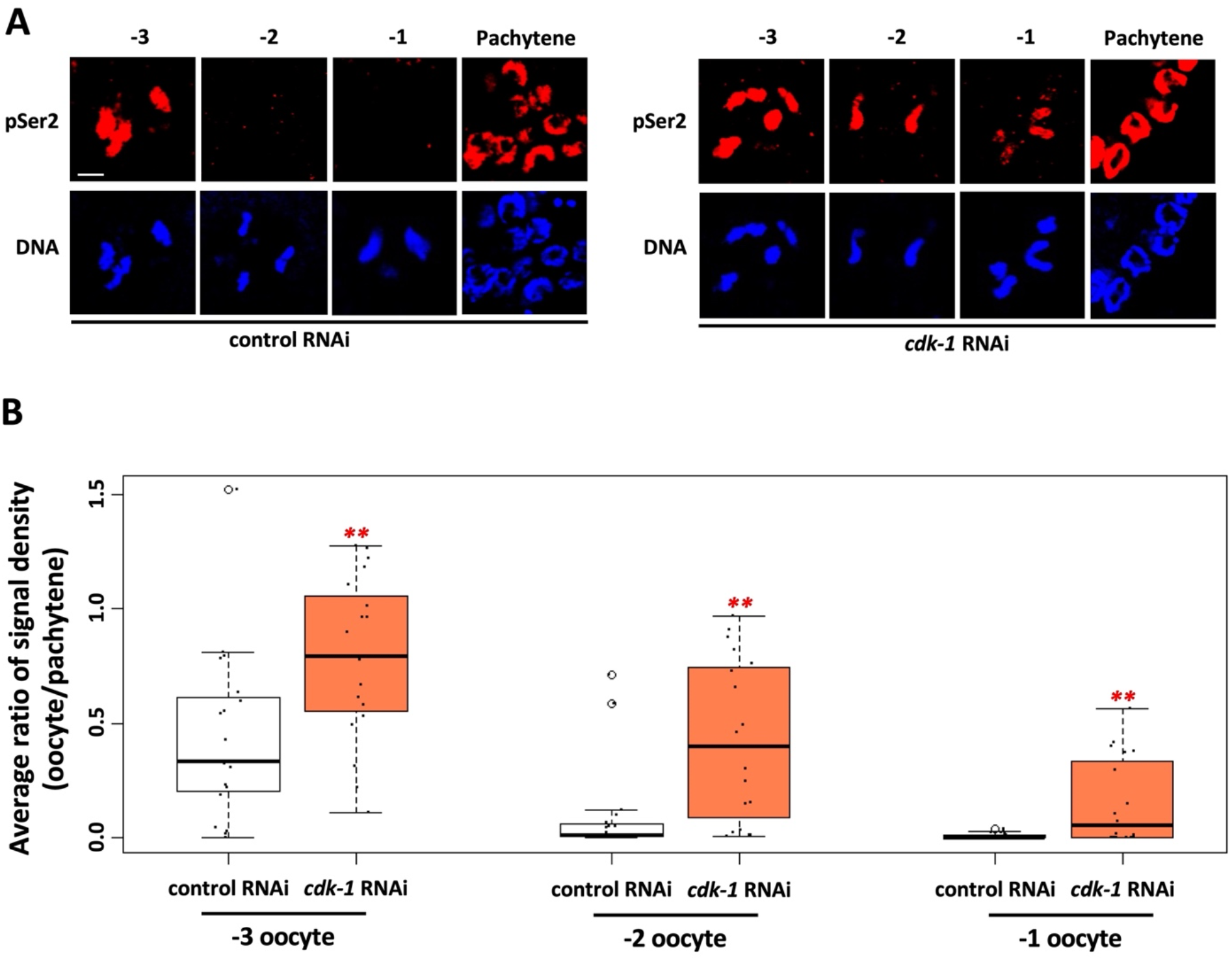
Aberrant RNAPIIpSer2 signal is observed in proximal oocytes when CDK-1 is depleted. A. N2 animals were treated with control or *cdk-1* RNAi. Gonads were dissected from these animals and were fixed and stained for DNA (blue) and RNAPIIpSer2 (red). Loss of CDK-1 results in RNAPIIpSer2 signal persisting into the −2 and −1 oocyte positions. B. Quantification of data presented in (A). RNAPIIpSer2 signal was measured using ImageJ. The average ratio of normalized raw integrated density is plotted on the y-axis. 19 samples were analyzed for each RNAi treatment over 2 independent replicates. Significance was measured using student’s t-test or Wilcoxon Rank Sum test. **p<0.01; *p<0.05. Depletion of CDK-1 resulted in a significant increase in on-chromatin RNAPIIpSer2 signal in the three most proximal oocytes.

**Figure 4:**
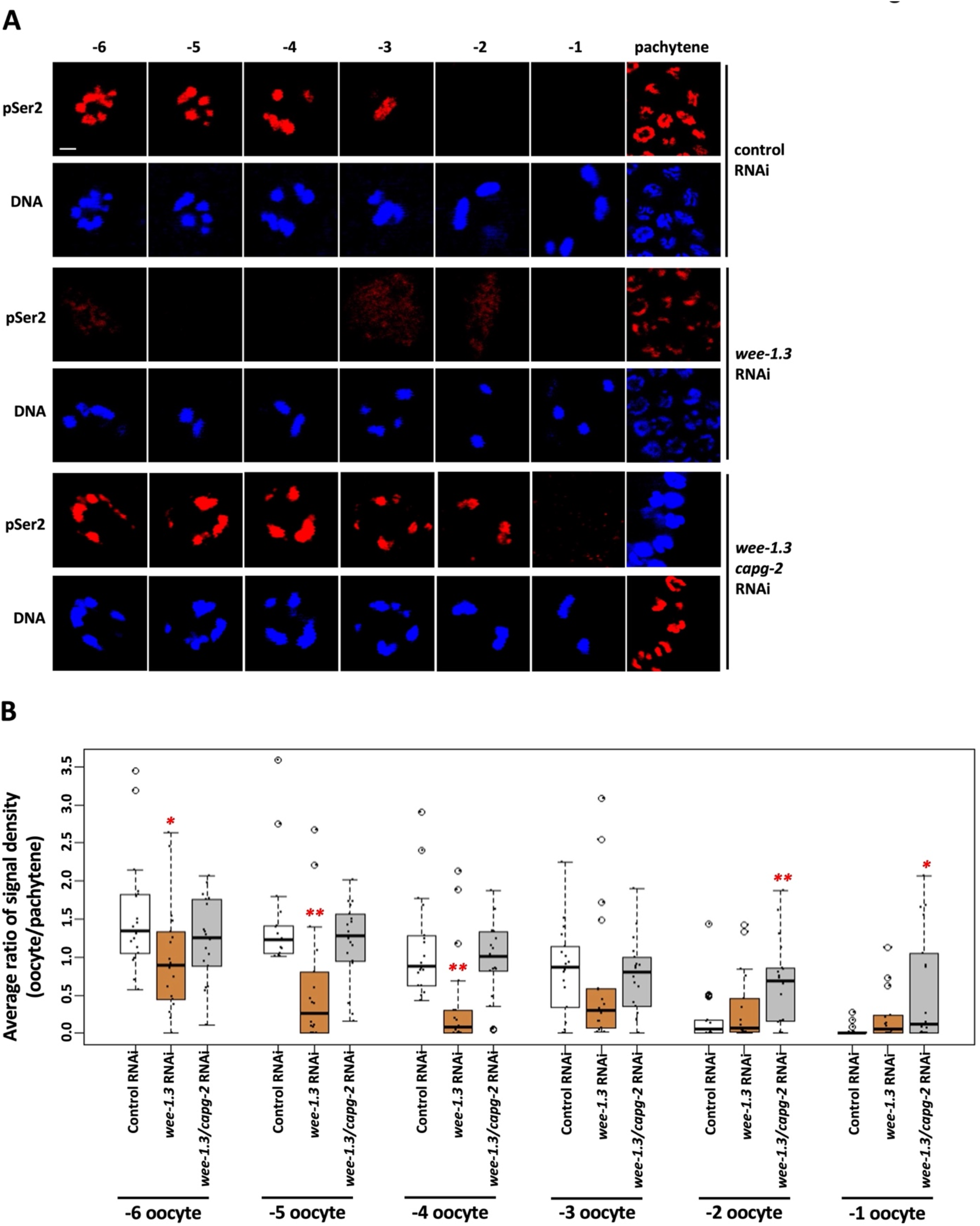
Hyperactivation of CDK-1 by the depletion of WEE-1.3 causes unscheduled transcriptional silencing. A. N2 animals were treated with either control, *wee-1.3*, or *wee-1.3/capg-2* double RNAi. Gonads were dissected from these animals and were fixed and stained for DNA (blue) and RNAPIIpSer2 (red). Depletion of WEE-1.3 results in the loss of on-chromatin RNAPIIpSer2 signals in the more distal (−6 to −4 position) oocytes. Co-depletion using *wee-1.3/capg-2* RNAi reverses the loss of RNAPIIpSer2 signal. Scale bars represent a length of 2 µm. B. Quantification of data presented in (A). RNAPIIpSer2 signal was measured using ImageJ. The average ratio of normalized raw integrated density is plotted on the y-axis. 20 samples were analyzed for each RNAi treatment over 2 independently performed replicates. Significance was measured using student’s t-test or Wilcoxon Rank Sum test. **p<0.01; *p<0.05. Depletion of WEE-1.3 resulted in a decrease in RNAPIIpSer2 signal from −6 to −2 oocyte positions. Co-depletion of WEE-1.3 and CAPG-2 resulted in restored RNAPIIpSer2 signal in −6 to −3 oocyte positions.

### H3K9me3 marks accumulate on oocyte chromatin during genome silencing

We next examined a role for the H3K9me pathway in oocyte genome silencing. Our previous work showed that the Z2/Z3 PGCs accumulate H3K9me marks to a significant extent, relative to neighboring somatic cells, as they prepare for silencing (Belew et al., 2021). To see if this also occurs on oocyte chromatin we used an antibody, termed ab176916 and purchased from Abcam (Waltham, MA), that recognizes the tri-methylated form of lysine 9 on histone H3. We have previously validated this antibody with the demonstration that reactivity is lost in Z2/Z3 of L1 larvae in a strain lacking SET-25, the major H3K9me3 methyltransferase in *C. elegans* (Belew et al., 2021). We limited the current analysis to H3K9me3 because available antibodies against H3K9me2 do not recognize their target when the neighboring serine 10 is phosphorylated (Kimura et al., 2008), and we have seen in Figure S2 that H3pS10 is prominent in proximal oocytes. As shown in Figure 5A, H3K9me3 first appears in −5 oocytes, and then gradually accumulates such that by the −2 and −1 positions the H3K9me3 and DNA signals overlap extensively. To examine the reproducibility of this pattern we quantified H3K9me3 signal intensity across 11 gonads and then compared the values obtained for the −5 to −2 positions to the value obtained for −1 position within a given gonad. As shown in Figure 5B, the pattern of gradual accumulation of H3K9me3 marks as oocytes moved from distal to proximal was indeed highly reproducible. In *C. elegans*, H3K9me3 is produced primarily by the SET-25 methyltransferase (Towbin et al., 2012). Indeed, when H3K9me3 was examined in *set-25* mutants and compared to wild type samples, we observed that the H3K9me3 signals were attenuated (Figure 5A). Figure 5C shows additional examples of −1 and −2 oocytes for wild type and *set-25* mutants at higher magnification, highlighting the differences in H3K9me3 signal intensity. No such attenuation was observed when *met-2* mutants were compared to wild type (Figure 5A), consistent with the MET-2 methyltransferase primarily responsible for producing H3K9me1 and me2 (Towbin et al., 2012). Our findings are also consistent with those of Bessler et al., 2010, who observed that H3K9me3 levels in the pachytene region of the gonad do not change in *met-2* mutants, relative to wild type. We conclude that H3K9me3 marks accumulate dramatically on chromatin at the time that oocytes are silencing their genomes.

**Figure 5:**
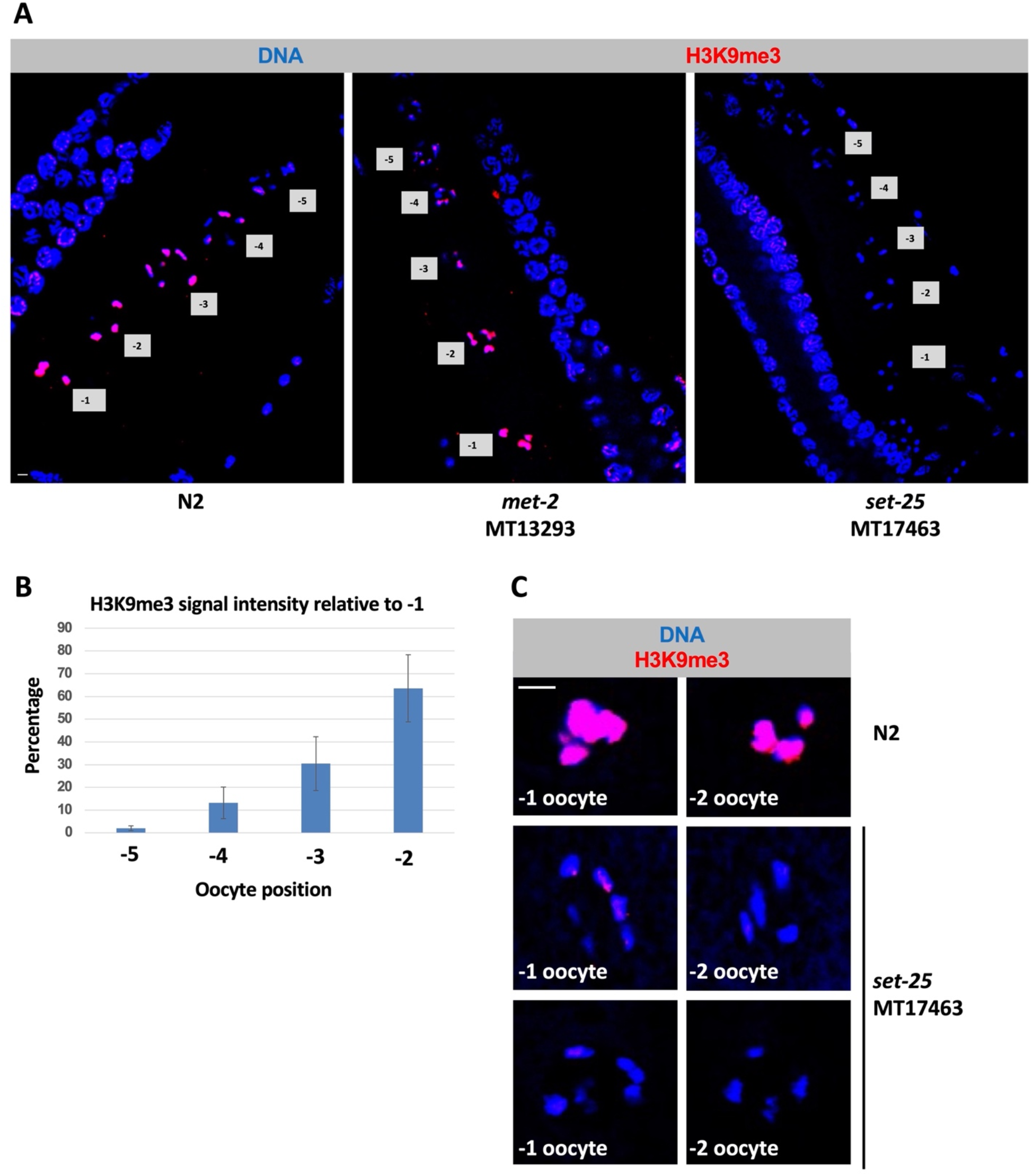
H3K9me3 signals significantly increase as oocytes become more proximal. A. Wild-type, *met-2* mutant, and *set-25* mutant gonads were dissected, fixed, and stained for DNA (blue) and H3K9me3 (red). In N2s, an increase in H3K9me3 signal is observed at the −3 to −1 positions in comparison to more distal oocytes and pachytene nuclei from the same animal. Loss of MET-2 does not affect the H3K9me3 accumulation pattern. Depletion of SET-25 results in loss of H3K9me3 signal in proximal oocytes. Scale bar represents a length of 2 µm. B. Quantification of data for wild type samples in (A). H3K9me3 signals for each oocyte in − 5 to −2 positions relative to the most proximal oocyte are presented. 20 samples were analyzed over 2 independent replicates. H3K9me3 signal accumulates on chromatin in the more proximal oocyte positions. C. Additional examples of −2 and −1 oocytes stained as in part (A). Either wild type (N2) or *set-25* mutant oocytes are shown. Note that the *set-25* samples have only trace amounts of H3K9me3 on the chromatin, relative to the wild type sample.

### Both the SET-25 and MET-2 methyltransferases are required for genome silencing in oocytes

Having observed SET-25 dependent accumulation of H3K9me3 marks in proximal oocytes we next asked if SET-25 plays a role in genome silencing. Staining of *set-25* mutants for RNAPIIpSer2 revealed that −2 oocytes contained significantly more RNAPIIpSer2 signal than did the control samples (Figure 6A&B), showing that silencing was attenuated. Interestingly, we also observed a silencing defect in *met-2* mutants (Figure 6A&B). Thus, not only do H3K9me marks accumulate during silencing, but the enzymes responsible for catalyzing these modifications are also important for genome silencing. We conclude that the H3K9me pathway globally silences transcription in developing oocytes, as it does in the PGCs of starved L1s.

**Figure 6:**
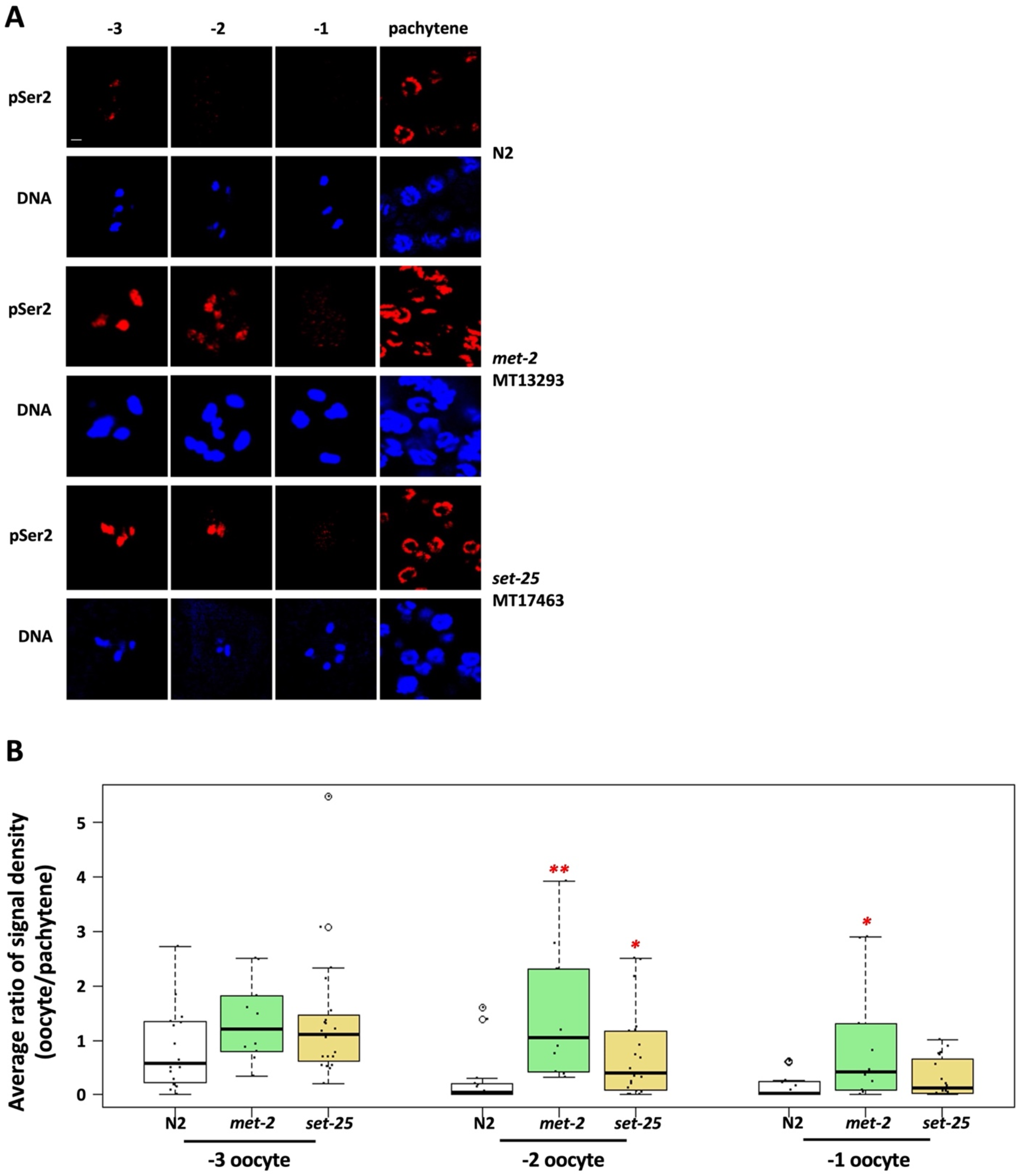
The H3K9 methyltransferases MET-2 and SET-25 are required for transcriptional repression in proximal oocytes. A. Gonads from wild-type, *met-2* mutant, and *set-25* mutant animals were dissected, fixed, and stained for DNA (blue) and RNAPIIpSer2 (red). Mutations of either *met-2* or *set-25* results in persistent RNAPIIpSer2 signal on chromatin, compared to wild-type animals. Scale bar represents a length of 2 µm. B. Quantification of data presented in (A). RNAPIIpSer2 signal was measured using ImageJ. The average ratio of normalized raw integrated density is plotted on the y-axis. 15 wild-type, 10 *met-2* mutants, and 20 *set-25* mutants were analyzed over 2 independent replicates. Significance was measured using student’s t-test or Wilcoxon Rank Sum test. **p<0.01; *p<0.05. Loss of MET-2 resulted in a significant persistence of RNAPIIpSer2 signal on chromatin in −2 and −1 oocytes. Loss of SET-25 led to a significant persistence of RNAPIIpSer2 signal on chromatin in −2 oocytes.

### Genome silencing is coupled to chromatin compaction in oocytes

Previous work has shown that condensin II loads on to oocyte chromosomes at the −3 position and is required for the intense chromatin compaction that occurs as bivalents are formed (Chan et al., 2004). In starved L1s, genome silencing is coupled to chromatin compaction (Belew et al., 2021), and thus it was important to monitor compaction in the oocyte system. For this we turned to a previously utilized strain that carries a transgene encoding mCherry-tagged histone H2B, which marks chromatin (Wong et al., 2018; Belew et al., 2021). Living hermaphrodites were immobilized and oocyte chromatin was imaged using confocal microscopy. We compared control samples to those that were exposed to *top-2/capg-2* RNAi, and we looked at −2 oocytes, which is where genome silencing is occurring. As described in the Methods and Figure S4, we measured the volume of the chromatin masses and found that depletion of TOP-2/CAPG-2 caused a significant increase in volume, consistent with a defect in compaction, and similar observations were made after *set-25* or *met-2* RNAi (Figures 7 and S4). Thus, as is the case in the PGCs of starved L1s, chromatin compaction in oocytes is driven by actions of the TOP-2/condensin II axis and components of the H3K9me pathway. We note that the effects observed here for bivalent compaction after *top-2/capg-*2 RNAi are less extreme than those reported by Chan and colleagues, and this is likely due to differences in how condensin II was inactivated (Chan et al., 2004). In our experiments, we use a feeding RNAi treatment that targets *capg-2*. By contrast, Chan and colleagues combined RNAi with a temperature-sensitive allele, both targeting the *hcp-6* subunit of condensin II, thereby using two forms of condensin II inactivation in the same experiment. As detailed by Chan and colleagues, the effect on bivalent compaction of the temperature-sensitive *hcp-6* (*hcp-6^ts^*) allele alone, without RNAi, is rather modest.

**Figure 7:**
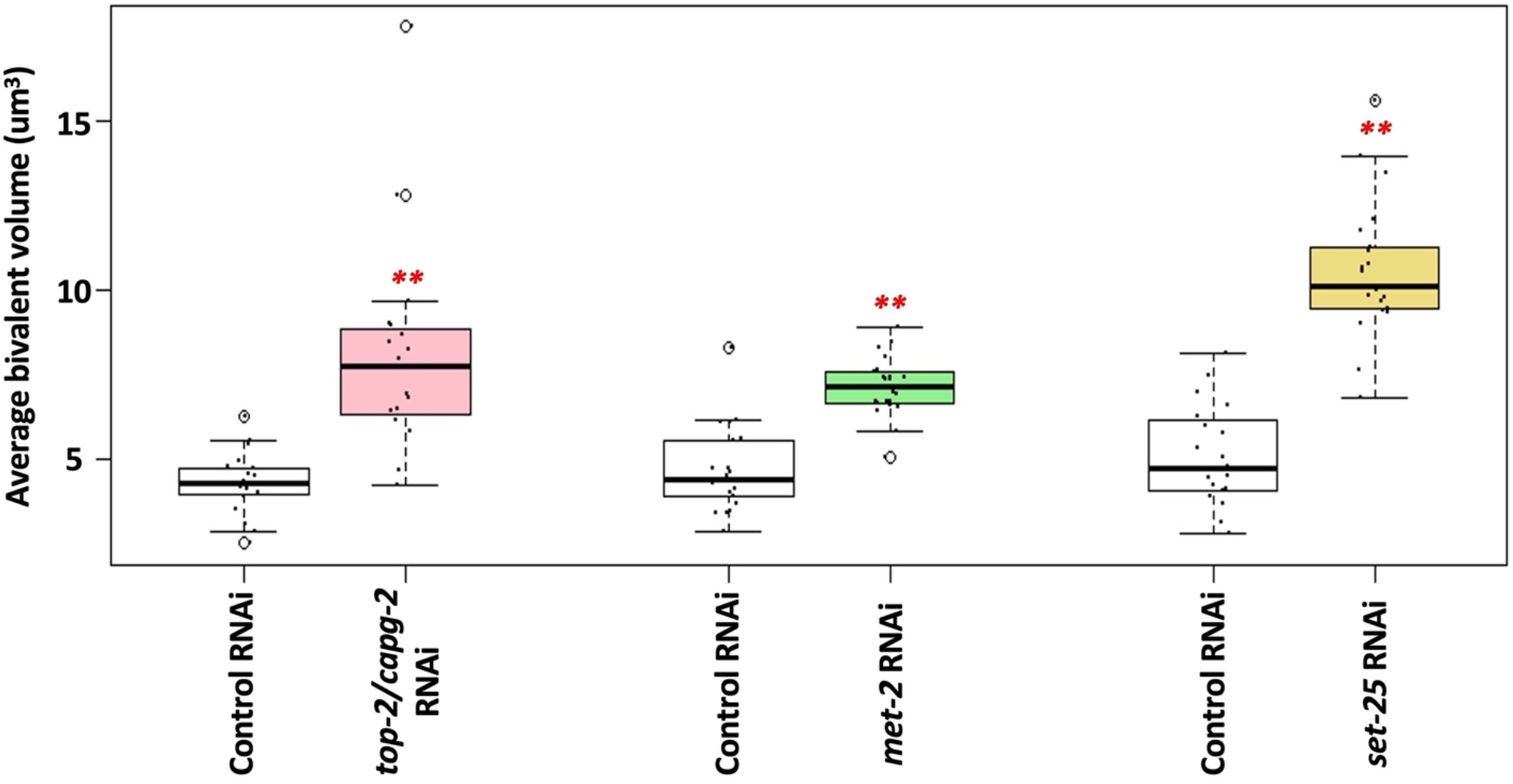
TOP-2, condensin II and the H3K9 methyltransferases MET-2 and SET-25 are all required for proper bivalent compaction. Living proximal oocytes, treated with control, *top-2/capg-2*, *met-2*, or *set-25* RNAi, were imaged for chromatin compaction using a strain harboring mCherry-tagged histone H2B. Bivalent volume in the −2 oocyte was measured using ImageJ. The average bivalent volume is plotted on the y-axis. 5 oocyte nuclei (with an average of 4 bivalents per nucleus) were analyzed for each RNAi treatment. Significance was measured using student’s t-test or Wilcoxon Rank Sum test. **p<0.01; *p<0.05. Exposure to *top-2/capg-2*, *met-2* or *set-25* RNAi treatments results in significantly larger bivalents.

### PIE-1 is required for genome silencing during meiotic prophase in oocytes and localizes to the nucleolus prior to silencing

In a final set of experiments, we examined a requirement for PIE-1 in oocyte silencing, as recent work has shown that PIE-1 is present in the adult gonad (Kim et al., 2021). Figure 8A shows that RNAi against *pie-1* causes a persistence of transcription in the −2 position, and quantification shows that RNAPIIpSer2 signal intensity is significantly higher in both −3 and −2 oocytes after *pie-1* RNAi, relative to the control samples (Figure 8B). Interestingly, PIE-1 depletion had no effect on bivalent compaction (Figure 8C). Thus, like TOP-2/condensin II and the H3K9me pathway, PIE-1 is required to repress transcription in −3/−2 oocytes, but unlike these factors its mechanism of action is distinct from chromatin compaction.

**Figure 8:**
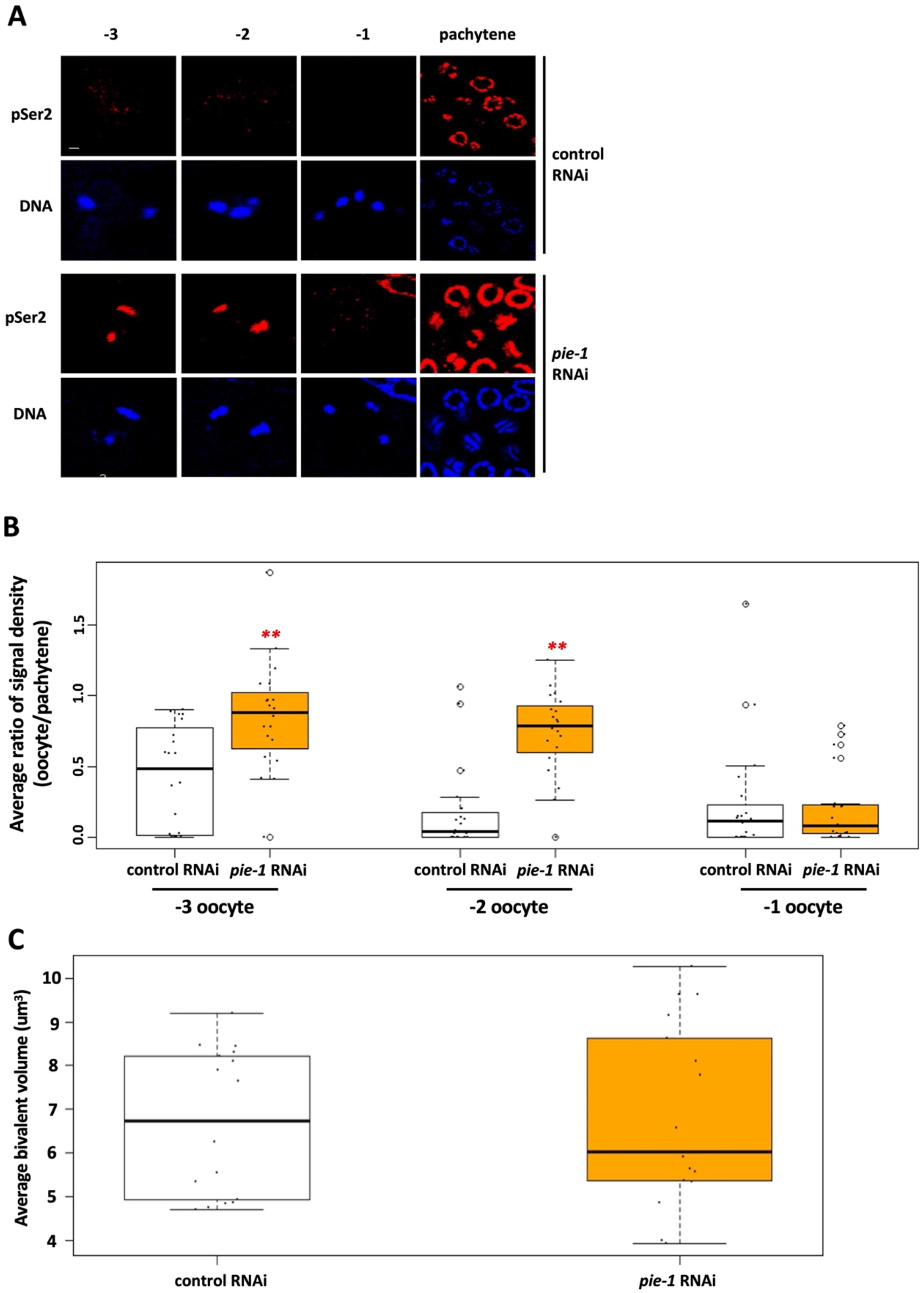
PIE-1 is required for transcriptional repression in proximal oocytes. A. N2 animals were treated with either control or *pie-1* RNAi. Gonads were dissected from these animals and were fixed and stained for DNA (blue) and RNAPIIpSer2 (red). Depletion of PIE-1 results in persisting RNAPIIpSer2 signal in proximal oocytes. B. Quantification of data presented in (A). RNAPIIpSer2 signal was measured using ImageJ. The average ratio of normalized raw integrated density is plotted on the y-axis. 20 samples were analyzed for each RNAi treatment over 2 independent replicates. Significance was measured using student’s t-test or Wilcoxon Rank Sum test. **p<0.01; *p<0.05. Oocytes from the *pie-1* RNAi treatment had significantly increased RNAPIIpSer2 signal at the −3 and −2 positions compared to control RNAi. C. Living proximal oocytes treated with control or *pie-1* RNAi were imaged for chromatin compaction using a strain harboring mCherry-tagged histone H2B. Bivalent volume in the −2 oocyte was measured using ImageJ. The average bivalent volume is plotted on the y-axis. 5 oocyte nuclei (with an average of 4 bivalents per nucleus) were analyzed for each RNAi treatment. Significance was measured using student’s t-test or Wilcoxon Rank Sum test. Exposure to *pie-1* RNAi treatment did not affect bivalent volume.

To pursue these findings, we next analyzed PIE-1 localization in oocytes, using a strain where GFP had been inserted at the endogenous *pie-1* locus (Kim et al., 2021). A typical localization pattern is shown in Figure 9A, where we see that PIE-1::GFP is mostly localized to the nucleus. Furthermore, within the −5 to −3 range of oocytes, it is clear that PIE-1::GFP accumulated within the nucleolus (Figures 9A&B), which can be easily observed using phase-contrast microscopy (Figure 9B). Previous work has shown that as oocytes prepare for maturation the nucleolus is lost, likely reflecting a shutdown of RNA polymerase I (RNAPI) transcription (Korčeková et al., 2012). This explains why PIE-1::GFP is no longer predominantly localized to the nucleolus in −2 oocytes, as the nucleolus is undergoing dissolution (Figures 9A&B). Previous work in budding yeast has shown that condensin is required to remodel rDNA chromatin in preparation for cell division (Freeman et al., 2000). Given this, we wondered if TOP-2/condensin II is required for nucleolar dissolution in proximal oocytes. To address this, we used phase-contrast microscopy to image nucleoli, and we simply measured their area in control and *top-2/capg-2 (RNAi)* samples. As shown in Figures 9 C&D, for control samples, we observed a significant decrease in nucleolar size in −2 oocytes, relative to −3 oocytes. This is consistent with the nucleolus undergoing dissolution in −2 oocytes. When nucleoli were assessed in samples depleted of TOP-2/CAPG-2, we saw that nucleolar size was significantly increased at both the −3 and −2 positions, relative to controls (Figures 9 C&D). Thus, the TOP-2/condensin II axis promotes nucleolar dissolution in proximal oocytes. We also imaged PIE-1::GFP in samples depleted of TOP-2/condensin II and observed, as expected, that PIE-1 was localized to the −2 nucleoli that had resisted dissolution (Figure 9E). These data show that the TOP-2/condensin II axis controls PIE-1::GFP localization, and this likely occurs via TOP-2/condensin II’s ability to promote nucleolar dissolution. As detailed below in the Discussion, these findings suggest a model for how TOP-2/condensin II and PIE-1 work together to promote genome silencing in proximal oocytes.

**Figure 9:**
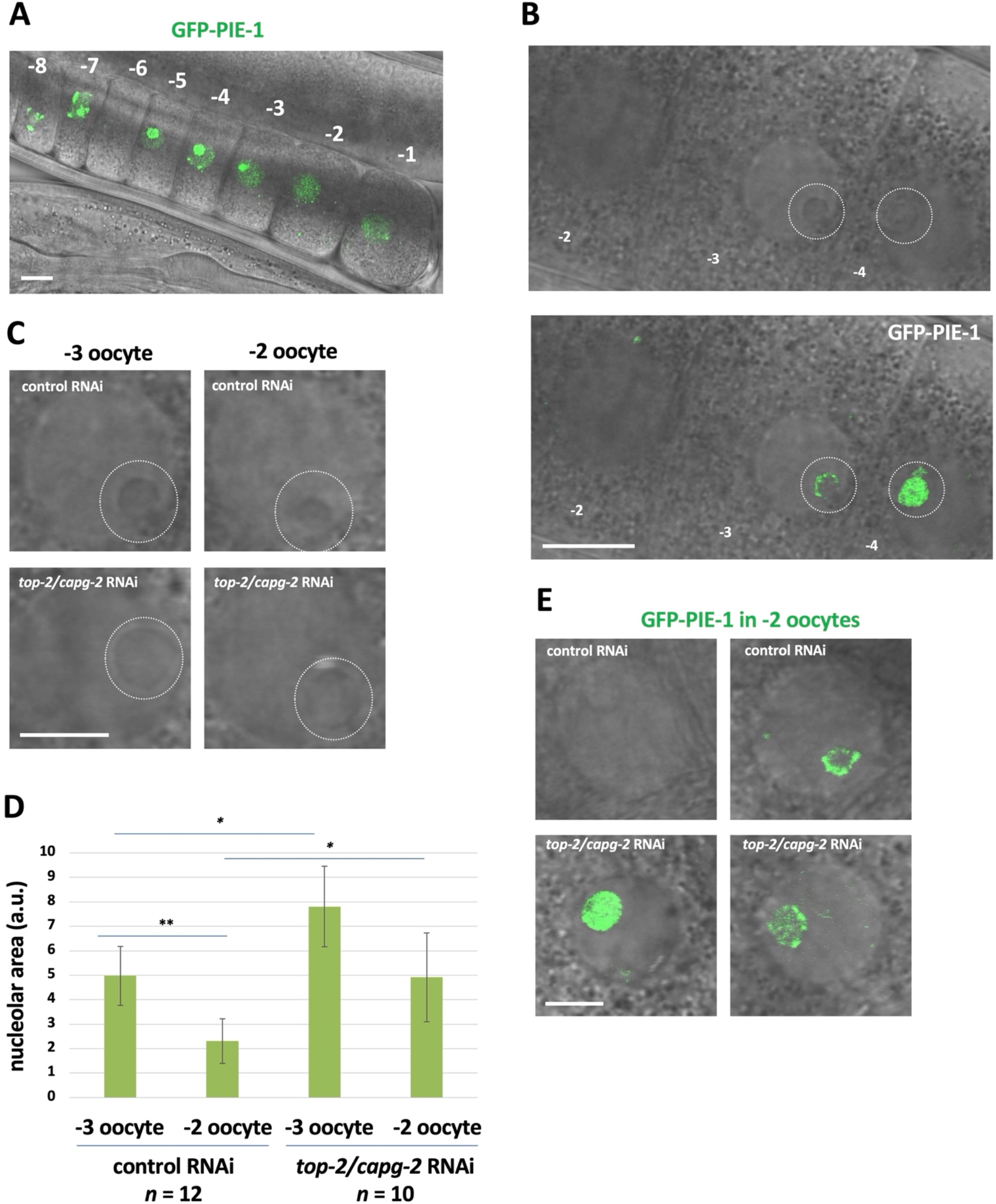
PIE-1 is sequestered in the nucleolus prior to silencing. A. The normal localization pattern of PIE-1::GFP in oocytes of live WM330 worms. From the −5 to −3 oocyte positions, PIE-1::GFP is sequestered within the nucleolus. Oocyte position is numbered. Scale bar represents a length of 10 µm. B. A wild-type localization pattern of PIE-1::GFP at the −3 oocyte position. PIE-1::GFP appears within the nucleolus. Nucleolus is indicated by the white circle. Scale bar represents a length of 10 µm. C. WM330 animals were treated with control or *top-2/capg-2* double RNAi. Live adults were imaged at the −3 and −2 oocyte positions. Scale bar represents a length of 5 µm. D. Quantification of data presented in (C). Nucleolar size was measured using ImageJ and is plotted on the y-axis (see Methods). Significance was measured using student’s t-test. **p<0.01; *p<0.05. Exposure to *top-2/capg-2* treatment results in significantly larger nucleoli. E. Live adults were imaged for PIE-1::GFP after treatment with control or *top-2/capg-2* double RNAi. Loss of *top-2/capg-*2 results in PIE-1::GFP remaining sequestered in the nucleolus at the −2 oocyte position. Scale bar represents a length of 5 µm.

## Discussion

The goal of this study was to define the molecular components for genome silencing in *C. elegans* oocytes. Unfortunately, there are no methods currently available to directly detect new mRNA transcription *in situ* in the worm gonad. Therefore, to monitor transcription, we relied on RNAPIIpSer2 staining, as we and others have done extensively in the past (see, for example, Seydoux and Dunn, 1997; Walker et al., 2007; Guven-Ozkan et al., 2008; Cassart and Yague-Sanz et al., 2020; Belew et al., 2021). One concern with this antibody-based approach is that chromatin compaction in −2 oocytes may prevent access of the antibody to its target on chromatin, rendering false-negative data. This is clearly not the case, however, as when we depleted PIE-1 we observed strong RNAPIIpSer2 signals on chromatin in −2 oocytes, even though compaction occurs normally under this condition (Figure 8). Thus, we consider RNAPIIpSer2 staining to be an accurate and legitimate method to assess the transcriptional status of oocytes in the worm.

Under normal conditions, we found that RNAPIIpSer2 signal intensity drops significantly in −2 oocytes relative to the −3 position, and that signals are often undetectable at −2 (Figure 2). This is consistent with previous work (Walker et al., 2007), and thus we conclude that genome silencing likely initiates at −3 and is largely complete by −2. At the −2 position, all oocytes have intact nuclear envelopes, and thus these cells have yet to enter meiotic M-phase. This is important as previous work has shown that transcription is repressed during M-phase (Taylor, 1960; Prescott and Bender, 1962; Parsons and Spencer; 1997), although more recent work has shown that the block is not absolute as a low-level of transcription can be detected in mitotic cells (Palozola et al., 2017). Nonetheless, it is clear that M-phase is incompatible with active transcription, and thus it is important to distinguish the genome silencing we observe in −2 oocytes from the repression of transcription that occurs during meiotic M-phase in −1 oocytes. Our data support this distinction in multiple ways. First, depletion of the genome silencing factors TOP-2, CAPG-2, MET-2, SET-25, and PIE-1 all yield a common phenotype – a persistence of RNAPIIpSer2 signal in −2 oocytes, and loss of the signal in −1 oocytes (Figures 2, 6, and 8). Thus, if loss of active transcription in −2 oocytes occurs mechanistically the same as in −1 oocytes, then we should see a persistence of signal in the −1 position also, and we clearly do not. Second, depletion of CDK-1 results in persistence of RNAPIIpSer2 signals in −2 <u>and</u> −1 oocytes (Figure 3). After exposure to *cdk-1* RNAi, −1 oocytes largely retain the nuclear envelope (data not shown), consistent with a failure to enter meiotic M-phase. This shows that it is not simply occupancy of the −1 position that represses transcription; rather, it is M-phase entry that does so. Taken together, these data show that genome silencing precedes entry into meiotic M-phase. Why has the worm evolved a system to block transcription at −2 when it is going to happen anyway at −1? We propose that active transcription may hamper the chromosomal remodeling that occurs at −2, and thus that genome silencing at −2 has evolved to allow proper bivalent formation.

Taking a candidate approach, we found that multiple genome silencing pathways are operational in −2 oocytes, as loss of the TOP-2/condensin II axis, the H3K9me pathway, and PIE-1 all impact silencing. We also found that silencing is under cell cycle control, as reducing CDK-1 activity prevents silencing in −2 oocytes and increasing CDK-1 activity promotes precocious silencing in oocytes distal to −2 (Figures 3 and 4). Furthermore, we obtained evidence that oocytes gradually approach the M-phase state, as MPM-2 antigens accumulate gradually in the proximal gonad, and not abruptly at the −1 position as one might expect if CDK-1 is abruptly activated in −1 oocytes (Figure S3). Taken together, these data paint a picture where CDK-1 activity increases gradually as a function of oocyte position in the proximal gonad, and that it is the −2 position where CDK-1 activity crosses a threshold sufficient to trigger silencing, but not yet sufficient to trigger entry into meiotic M-phase. Thus, we propose that the timing of genome silencing is controlled by a CDK-1 activity gradient spanning the proximal gonad (Figure 10A).

**Figure 10:**
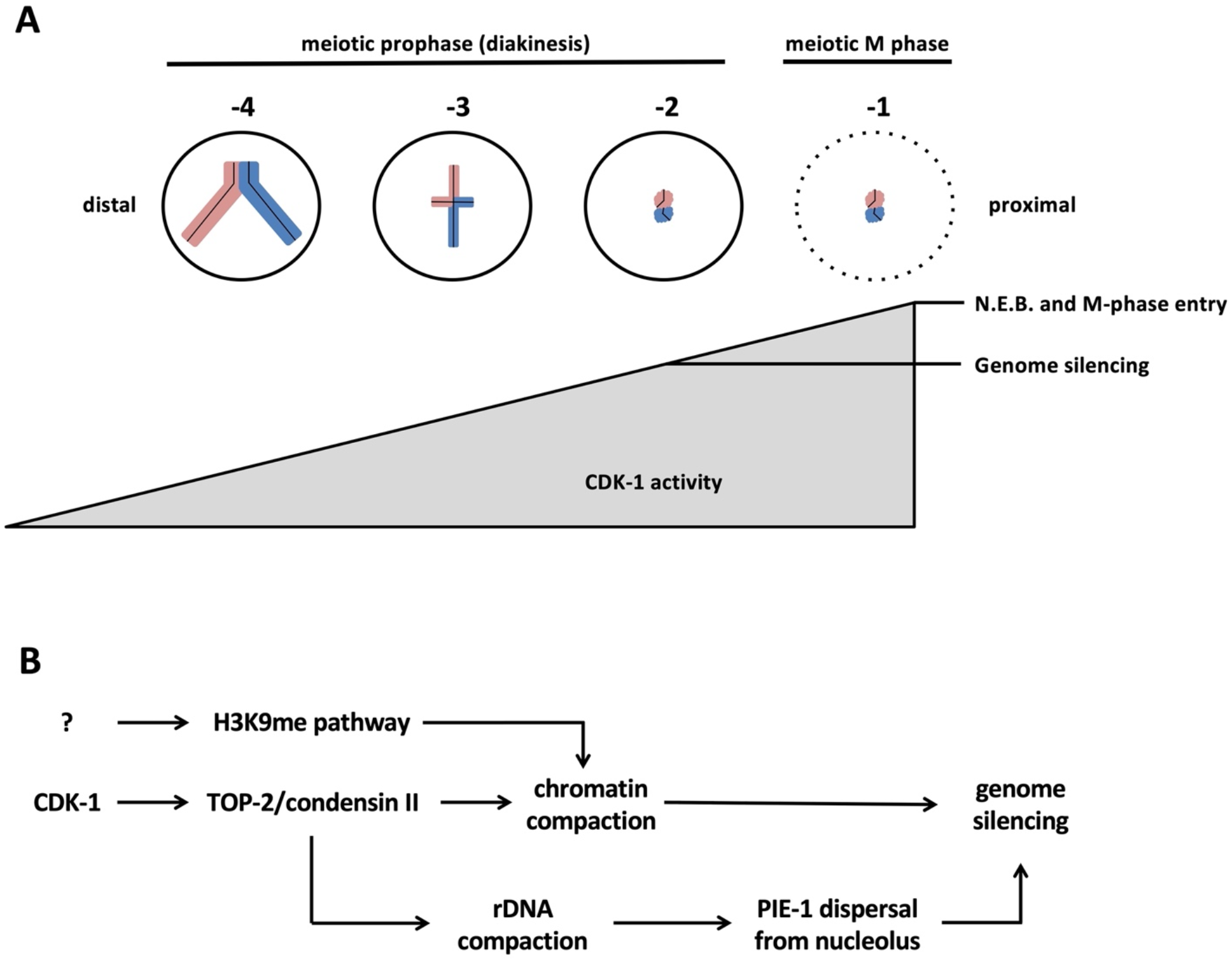
Models for how genome silencing occurs in proximal oocytes. A. A CDK-1 activity gradient allows for genome silencing at the −2 position to occur prior to entry into meiotic M-phase at the −1 position. B. A proposed pathway whereby CDK-1 promotes TOP-2/condensin II mediated compaction of rDNA, and this in turn promotes nucleolar dissolution and relocalization of PIE-1 to the nucleoplasm where it can block transcription.

Our previous work had identified a linear pathway, termed GCC, that silences germline transcription during L1 starvation (Belew et al., 2021). For GCC, TOP-2/condensin II acts upstream of the MET-2 and SET-25 methyltransferases to promote H3K9me2 and -me3 deposition on chromatin (Belew et al., 2021). Interestingly, while TOP-2/condensin II and SET-25/MET-2 are all required for silencing in oocytes, they are not organized into a linear pathway as we found that H3K9me3 deposition in oocytes does not require TOP-2/condensin II (data not shown). H3K9me3 deposition is increased as oocytes approach the −1 position (Figure 5), and this is similar to what happens in late embryos, where deposition is dramatically increased in Z2/Z3, relative to their somatic counterparts (Belew et al., 2021). How this increase is regulated, and whether it occurs via spreading of preexisting marks or via *de novo* deposition are fascinating questions for future research. As mentioned above, we were unable to track H3K9me2 marks, but it is interesting to note that loss of *met-2*, which is responsible for H3K9me2 deposition, has a stronger effect on silencing than does loss of *set-25*, which is responsible for H3K9me3 (Figure 6). It thus appears that both H3K9me2 and -me3 play a role in genome silencing, and another important route for future research will be to determine which of the H3K9me readers is involved and how they are acting mechanistically to promote silencing. We note that in mice, it has been appreciated for some time that oocytes undergo chromatin compaction concurrent with transcriptional repression (Zuccotti et al., 1995; Bouniol-Baly et al., 1999; De La Fuente et al., 2004; Schultz et al., 2018). Interestingly, very recent work has shown that a histone H3.3 chaperone complex, comprised of the Hira and Cabin1 proteins, promotes H3K9me3 deposition in chromatin, chromatin compaction, and genome silencing in mouse oocytes (Smith et al., 2022). These data are consistent with our findings and suggest that a conserved feature of genome silencing is the H3K9me-mediated compaction of chromatin on a global scale. It will be of interest to determine if TOP-2 and/or condensin II are also required for genome silencing in murine oocytes.

Our work also describes a novel function for PIE-1 in oocyte genome silencing. PIE-1 has been well studied in its role of blocking transcription in the P-lineage of early embryos, however how it does so is still a mystery. Early work suggested a model where PIE-1 binds to and sequesters cyclin T, a subunit of the CDK9 kinase that phosphorylates RNAPII on serine 2 within the CTD (Zhang et al., 2003). More recent work, however, has shown that PIE-1 mutants that fail to bind cyclin T still block transcription in the P-lineage (Ghosh and Seydoux, 2008) and, furthermore, that the RNAPIIpSer2 mark itself is dispensable for embryogenesis (Cassart and Yague-Sanz et al., 2020). Thus, it may be that, in the embryo, PIE-1 acts by interfering with RNAPII serine 5 phosphorylation (Ghosh and Seydoux, 2008), via an unknown mechanism. If so, this begs the question of why has PIE-1 evolved the ability to specifically interact with cyclin T? One plausible answer is that PIE-1 targets cyclin T to block transcription in oocytes. Interestingly, we have shown here that CDK-9, presumably acting with cyclin T, is the relevant serine 2 kinase in proximal oocytes, unlike the remainder of the gonad where CDK-12 is the relevant kinase (Bowman et al., 2013). This finding makes it intriguing to speculate that the switch from CDK-12 to CDK-9 occurs so that RNAPIIpSer2 can be regulated by PIE-1 in proximal oocytes. Sorting out how PIE-1 is blocking transcription in oocytes is another important avenue for future research.

Lastly, another important research question raised by our study is how is PIE-1 regulated in the proximal gonad? We have shown that PIE-1-GFP is present in the nuclei of −5, −4, and −3 oocytes, yet these cells are transcriptionally active (Figure 1B and 9A). Importantly, in oocytes distal to −2, PIE-1-GFP is localized predominantly in nucleoli (Figure 9). This might explain why these nuclei are competent for transcription despite the presence of PIE-1, if nucleolar residency prevents PIE-1 from accessing its target(s) for transcriptional repression. Previous work has shown that the nucleolus dissolves at the −2 position (Korčeková et al., 2012). We have observed this as well and, furthermore, we have shown that dissolution requires TOP-2/condensin II (Figure 9). How might the various components required for genome silencing in oocytes fit together mechanistically? Our data support the model shown in Figure 10B. We propose that once CDK-1 activity passes a threshold then TOP-2/condensin II is activated and recruited to chromatin, and this has two consequences. One, chromatin compaction commences and this represses transcription, likely via occlusion of RNAPII and various transcription factors from promoters on the compacted chromatin. Two, as the rDNA is compacted, RNAPI synthesis is blocked and the nucleolus dissolves, thereby liberating PIE-1 to block transcription via an unknown mechanism. Lastly, independent of TOP-2/condensin II, the methyltransferases targeting H3K9 are stimulated and H3K9me deposition is hyper-activated, leading to chromatin compaction and genome silencing. While this model is consistent with our data, there is clearly much more work needed to establish its accuracy. Future experiments will address the role of PIE-1 nucleolar residency in its regulation as well as the mechanism by which the SET-25 and MET-2 methyltransferases are activated and how H3K9me marks accumulate so dramatically on oocyte chromatin.

## Materials and Methods

### C. elegans strains

N2 (wild-type), WMM1 ([pie-1::gfp::pgl-1 + unc-119(+)]; [(pAA64)pie-1p::mCherry::his-58 + unc-119(+)] IV), MT13293(met-2(n4256) III), MT17463 (set-25(n5021) III) and WM330 (pie-1(ne4301[pie-1::GFP]) III) strains were used in this study. Worms were maintained on 60mm plates containing nematode growth media (NGM) seeded with the *E. coli* strain OP50 or HT115. Worms were grown at 20°C and propagated through egg preparation (bleaching) every 72 hours.

### Bacterial strains

OP50 bacteria served as the primary food source. It was grown in LB media containing 100 μg/ml streptomycin by shaking at 37°C overnight. 500 μl of the culture was seeded on Petri dishes containing NGM + streptomycin. HT115 bacteria grown in LB media containing 100 μg/ml carbenicillin and 12.5 μg/ml tetracycline and seeded on NGM + carbenicillin + tetracycline plates were also used as a source of food. Our RNAi strains were obtained from the Ahringer library and verified by Sanger sequencing. Bacteria containing dsRNA were streaked on LB-agar plates containing 100 μg/ml carbenicillin and 12.5 μg/ml tetracycline and incubated at 37°C overnight. Single colonies were then picked and grown in 25 ml LB cultures with 100 μg/ml carbenicillin and 12.5 μg/ml tetracycline. 500 μl of this culture was seeded on 60-mm Petri dishes containing 5mM IPTG.

### Egg preparation

Bleach solution containing 3.675 ml H_2_O, 1.2 NaOCl, and 0.125 ml 10N NaOH was prepared. Adult worms were washed from plates with 5 ml of M9 minimal medium (22mM KH_2_PO_4_, 22mM Na_2_HPO_4_, 85mM NaCl, and 2mM MgSO_4_). Worms were centrifuged at 1.9 KRPM for 1 minute and the excess medium was removed, then the bleach solution was added. Eggs were extracted by vortexing for 30 seconds and shaking for 1 minute. This was done a total of 3 times and worms were vortexed one last time. Then the eggs were spun down at 1900 rpm for 1 minute and excess bleach solution was removed, and the eggs were washed 3 times with M9 minimal medium.

### RNAi treatment

RNAi containing NGM plates were prepared as described in the “Bacterial strains” section. For double RNAi treatments, RNAi cultures were mixed at a 1:1 ratio by volume. HT115 cells transformed with an empty pL4440 vector was used as a negative control. RNAi conditions used in this study and tests for their efficacy is described below:

#### *cdk-9* RNAi

L1 worms were plated on HT115 food plates for the first 48 hours and were then moved to plates containing *cdk-9* RNAi for the remaining 24 hours. Embryonic lethality in the range of 80% - 85% was observed.

#### *top-2* RNAi

L1 worms were plated on HT115 food plates for the first 24 hours and were then moved to plates seeded with *top-2* RNAi for the remaining 48 hours. Embryonic lethality was observed at >90%. *<u>capg-2</u>* <u>RNAi</u> Worms were grown on HT115 food plates for the first 24 hours and were moved to plates containing *capg-2* RNAi for the remaining 48 hours. An embryonic lethality of 80%-100% was seen with this RNAi treatment.

#### *top-2/capg-2* double RNAi

Worms were grown on HT115 food plates for the first 24 hours and were transferred to *top-2/capg-2* double RNAi plates for the next 48 hours. Embryonic lethality ranged from 90%-100% for this RNAi treatment.

#### *cdk-1* RNAi

Worms were grown on HT115 food plates for the first 24 hours and were transferred to *cdk-1* RNAi plates for the next 48 hours. Embryonic lethality ranged from 97%-100% for this RNAi treatment.

#### *wee-1.3* RNAi

Worms were grown on plates containing *wee-1.3* RNAi for the entirety of their life cycle. An embryonic lethality of approximately 40% was observed. Additionally, a significant reduction in brood size, and the coalescence of bivalents into one chromatin mass in proximal oocytes of some samples, were observed for *wee-1.3* RNAi worms, as previously reported (Burrows et al., 2006).

#### *wee-1.3/capg-2* double RNAi

Worms were grown on plates containing *wee-1.3* RNAi for 24 hours then were moved to plates containing *wee-1.3/capg-2* RNAi where they remained for the rest of their life cycle. An embryonic lethality of approximately80%, and the coalescence of bivalents in proximal oocytes of some samples were observed.

#### *met-2* RNAi

Worms were grown on plates containing *met-2* RNAi for the entirety of their life cycle. Some of the adult worms were bleached and an L1 chromatin compaction assay was performed on the resulting larvae to test RNAi efficacy. See Belew et al., 2021, for details on the L1 compaction assay.

#### *set-25* RNAi

Worms were grown on plates containing *set-25* RNAi for the entirety of their life cycle. RNAi efficacy was tested via the same method as for *met-2* RNAi.

#### *pie-1* RNAi

Worms were grown on *pie-1* RNAi plates for the entirety of their life cycle. An embryonic lethality of 100% was observed for this RNAi.

### Antibodies and dilutions

<u>RNAPIIpSer2</u>: Rabbit antibody from Abcam (ab5095, Waltham, Massachusetts) was used at a dilution of 1:100. <u>H3pSer10</u>: Rabbit antibody from Rockland Immunochemicals (600-401-I74, Pottstown, Pennsylvania) was used at a dilution of 1:500. <u>H3K9me3</u>: Rabbit antibody from Abcam (ab176916, Waltham, Massachusetts) was used at a dilution of 1:1000. <u>MPM-2</u>: Mouse antibody (isotype - IgG1) from Sigma-Aldrich (05-368, St. Louis, Missouri) was used at a dilution of 1:500. <u>Secondary antibodies</u>: Alexa Fluor conjugated secondary antibodies from Invitrogen (Thermo Fisher Scientific, Waltham, Massachusetts) were used at a dilution of 1:200.

### Immunofluorescence staining

Adult worms were first washed off plates with 10 ml of M9 minimal medium and rinsed 3 more times. Then, they were centrifuged at 1.9 KRPM and the excess medium was removed. 20 μl of media containing about 50 worms were spotted on a coverslip and 3 μl of anesthetic (20mM Sodium Azide and 0.8M Tetramisole hydrochloride) was added to immobilize them. Worms were dissected using 25Gx5/8 needles (Sigma Aldrich, St. Louis, Missouri). To release gonads, adult worms were cut twice, once right below their pharyngeal bulb and once near the tail. The coverslip was then mounted onto poly-L-lysine covered slides and let rest for 5 minutes. Slides were put on dry ice for 30 minutes. Samples were then freeze-cracked by flicking the coverslips off for permeabilization.

For RNAPIIpSer2, H3pSer10, and MPM-2 antibody staining experiments, once samples were permeabilized, slides were put in cold 100% methanol (−20°C) for 2 minutes and then fixing solution (0.08M HEPES pH 6.9, 1.6mM MgSO_4_, 0.8mM EGTA, 3.7% formaldehyde, 1X phosphate-buffered saline) for another 30 minutes. After fixing, slides were washed three times with TBS-T (TBS with 0.1% Tween-20) and were blocked for 30 minutes with TNB (containing 100mM Tris-HCl, 200 mM NaCl, and 1% BSA). Primary antibodies were then applied at the dilutions described above in TNB and slides were incubated at 4°C overnight.

For H3K9me3 staining experiments, permeabilized samples were put in cold 100% methanol (−20°C) for 10 seconds and then fixing solution (0.08M HEPES pH 6.9, 1.6mM MgSO_4_, 0.8mM EGTA, 3.7% formaldehyde, 1X phosphate-buffered saline) for 10 minutes. After fixing, slides were washed three times with TBS-T (TBS with 0.1% Tween-20) and were blocked for 2 hours with TNB (containing 100mM Tris-HCl, 200 mM NaCl, and 1% BSA) supplemented with 10% normal goat serum. Primary antibodies were then applied at the dilutions described above in TNB supplemented with 10 % goat serum and slides were incubated at 4°C overnight.

On the next day, the slides were washed 3 times with TBS and slides were incubated with secondary antibodies and Hoechst 33342 dye for 2 hours at room temperature. Slides were washed 3 times with TBS, mounting medium (50 % glycerol in PBS), and coverslips were applied and sealed with Cytoseal XYL (Thermo Fisher Scientific, Waltham Massachusetts).

### Live animal imaging

Adult WMM1 and EGW83 worms were collected off plates and were washed 3 times with 10 ml M9 minimal medium. After the last wash, worms were spun down at 1.9 KRPM and excess medium was removed. 0.3% agarose pads were made on slides, and a 10 μl aliquot of adult worms was mounted. 4 μl of anesthetic (20mM Sodium Azide and 0.8M Tetramisole hydrochloride) was added to induce paralysis . A coverslip was gently applied, and the slides were imaged.

### Immunofluorescent imaging

All slides were imaged using an Olympus Fluoview FV1000 confocal microscope using Fluoview Viewer software at a magnification of 600x (60x objective and 10x eyepiece magnifications). Laser intensity was controlled for experiments to achieve consistency among samples.

### Quantification of data

#### RNAPIIpSer2 signal quantification

For each oocyte nucleus, two images were taken: one of the Hoechst-stained DNA and one of the RNAPIIpSer2 signal. Images were analyzed using ImageJ. An outline was drawn using the polygon selection tool around the Hoechst-stained area to mark the space occupied by DNA. The region of interest was copied and pasted to the RNAPIIpSer2 signal image and the raw integrated density (the sum of the values of the pixels in the selection) was measured. The raw integrated density was then normalized by the area to get a measure we called “signal density”. To account for possible variability in signal intensity due to different degrees of antibody penetration from sample to sample, each oocyte’s signal density was normalized to an average signal density from 5 pachytene nuclei found in the same image. The final normalized signal densities for the oocytes are presented. For each condition, data were collected from two independently performed experimental replicates and the data were then pooled for statistical analysis and presentation.

#### H3K9me3 signal quantification

Signal densities for each oocyte were calculated similarly to what is described above in the RNAPIIpSer2 signal quantification. H3K9me3 signal of each oocyte in the −2 to −5 positions was normalized to the most proximal (−1) oocyte and the ratios are presented as percentages.

#### H3pSer10 signal quantification

Signal densities for each oocyte were calculated similarly to what is described above in the RNAPIIpSer2 signal quantification. Signal densities were not normalized to pachytene nuclei since pachytene nuclei either did not harbor any signal (for H3pSer10 staining) or the signals were affected by our RNAi treatments (for H3K9me3 staining) which precluded us from using them for unbiased normalization.

#### Bivalent volume quantification

Z-stacks of oocyte nuclei were acquired. For each condition, 5 nuclei were analyzed and every bivalent within those nuclei that were visually distinguishable from one another was included in our analysis. On average 4 bivalents per nucleus were analyzed using ImageJ. The scale was set to 69nm per pixel. To measure the volume of a bivalent, a polygon was tightly drawn around it on each stack the bivalent appears in. The areas of these polygons were measured and summed up. Finally, the sum was multiplied by the distance between each stack to calculate an approximation of the bivalent’s volume. Averages of the volumes of all bivalents analyzed were presented.

#### Nucleolar size measurement

Images were captured of focal planes corresponding to the maximum diameter of the nucleolus in living oocytes for each oocyte position. Diameters were measured using ImageJ and then converted to area.

### Statistical analysis

Prior to performing any statistical test, data was tested whether it was parametric or not. To do so, the Shapiro-Wilk test was used to test for normal distribution and F-test was used to test for variance homogeneity of the datasets we were comparing. Data were then analyzed using a student’s t-test or Wilcoxon Rank Sum test depending on whether the datasets fulfill the requirements for a parametric test or not. Differences between any two datasets were considered statistically significant if a P-value of <0.05 was obtained.

## Author Contributions

W.M.M. acquired funding and conceptualized the project. W.M.M designed experiments. M.D.B., E.C., M.M.W., and W.M.M. performed experiments. W.M.M., M.D.B., and E.C. wrote the manuscript.

## Acknowledgements

We are grateful to Erik Griffin and Craig Mello for the kind gifts of worm strains.

---

**Figure S1:**
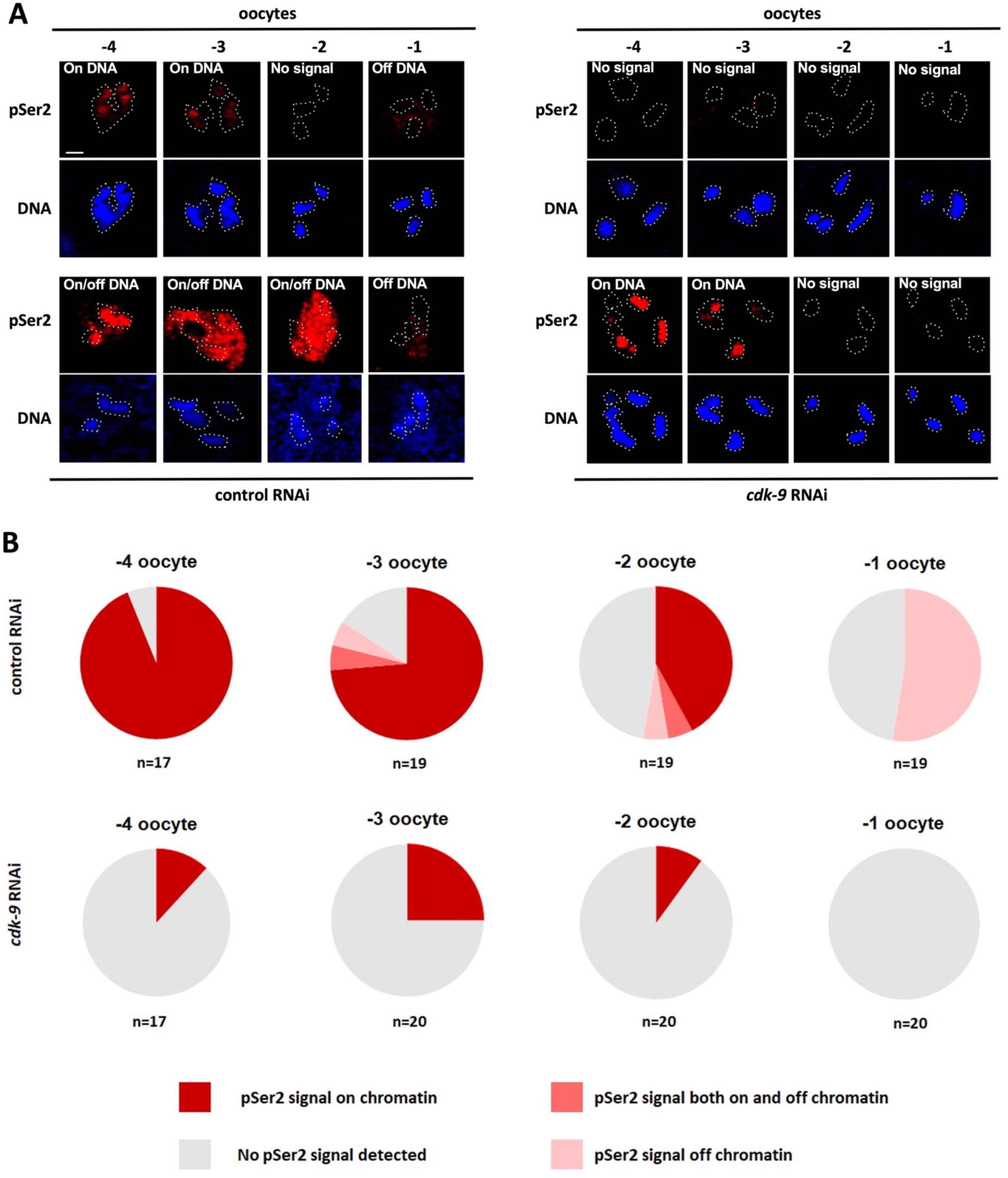
All patterns of RNAPIIpSer2 signal in *C. elegans* proximal oocytes are dependent on CDK-9. A. The different patterns of RNAPIIpSer2 (red) signal on and/or off DNA (blue) observed in N2s treated either control or *cdk-9* RNAi. Depletion of CDK-9 results in the loss of RNAPIIpSer2 signal in proximal oocytes regardless of RNAPIIpSer2 signal localization. Scale bar represents a length of 2 µm. B. Visualization of data presented in (A). N2s treated with *cdk-9* RNAi showed reduced RNAPIIpSer2 signal in all forms. The number of samples analyzed over 2 independent replicates is presented below the charts for each oocyte position.

**Figure S2:**
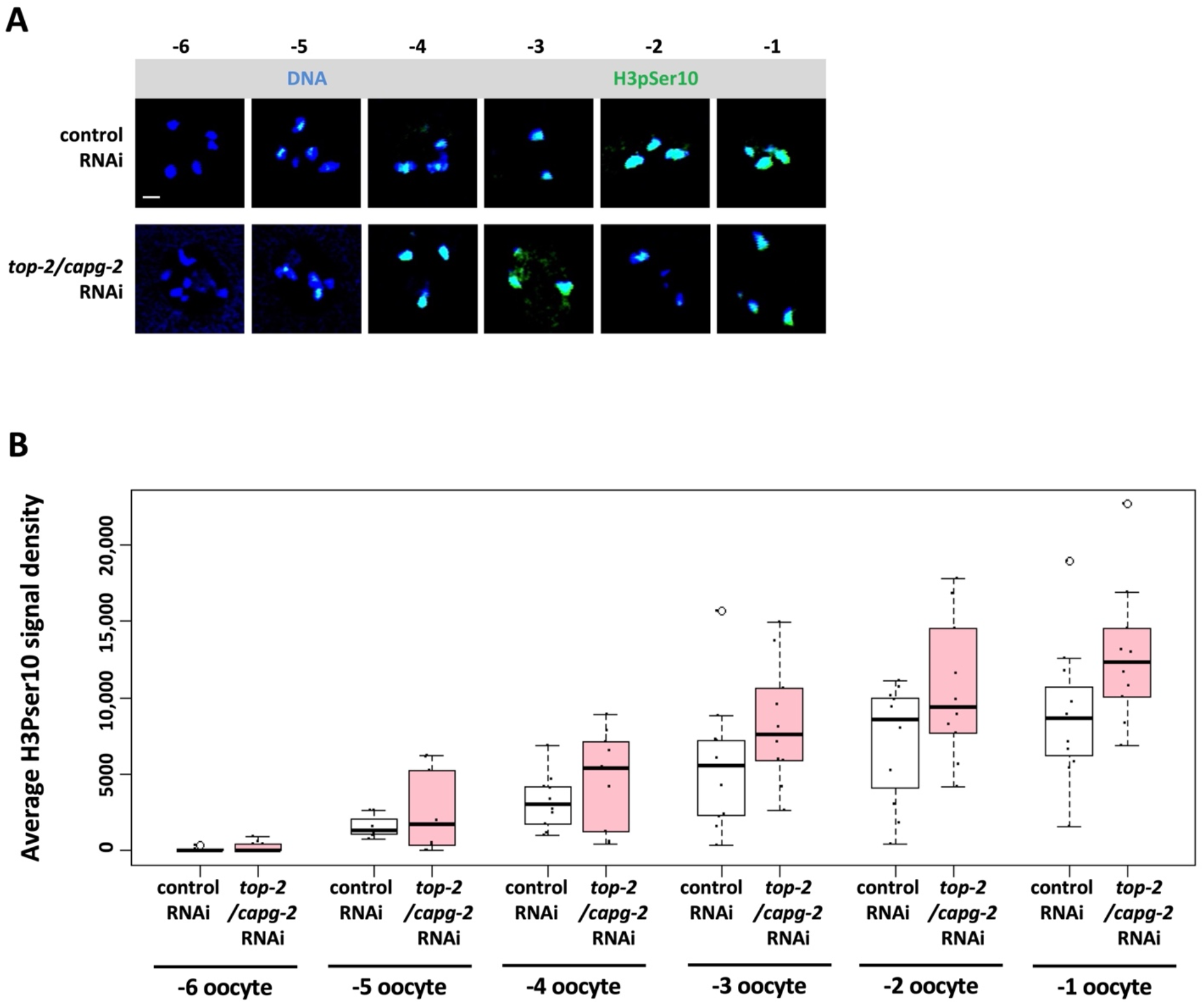
TOP-2 and condensin II mediated transcriptional repression in oocytes is independent of cell-cycle timing. A. N2 animals were treated with control or *top-2/capg-2* double RNAi. Gonads from young adults were dissected, fixed, and stained for DNA (blue) and H3pSer10 (green). Exposure to *top-2/capg-2* RNAi does not affect the timing of H3pSer10. Scale bar represents a length of 2 µm. B. Quantification of data presented in (A). 11 control RNAi and 10 *top-2/capg-2* RNAi samples were analyzed for each RNAi treatment. The onset of H3pSer10 occurs at the −5 position in both RNAi conditions, showing that the timing of prophase progression is not altered after *top-2/capg-2* RNAi. Signal density was not normalized due to the lack of an appropriate control tissue, and thus the basis for the difference in H3pSer10 intensity is presently unclear.

**Figure S3:**
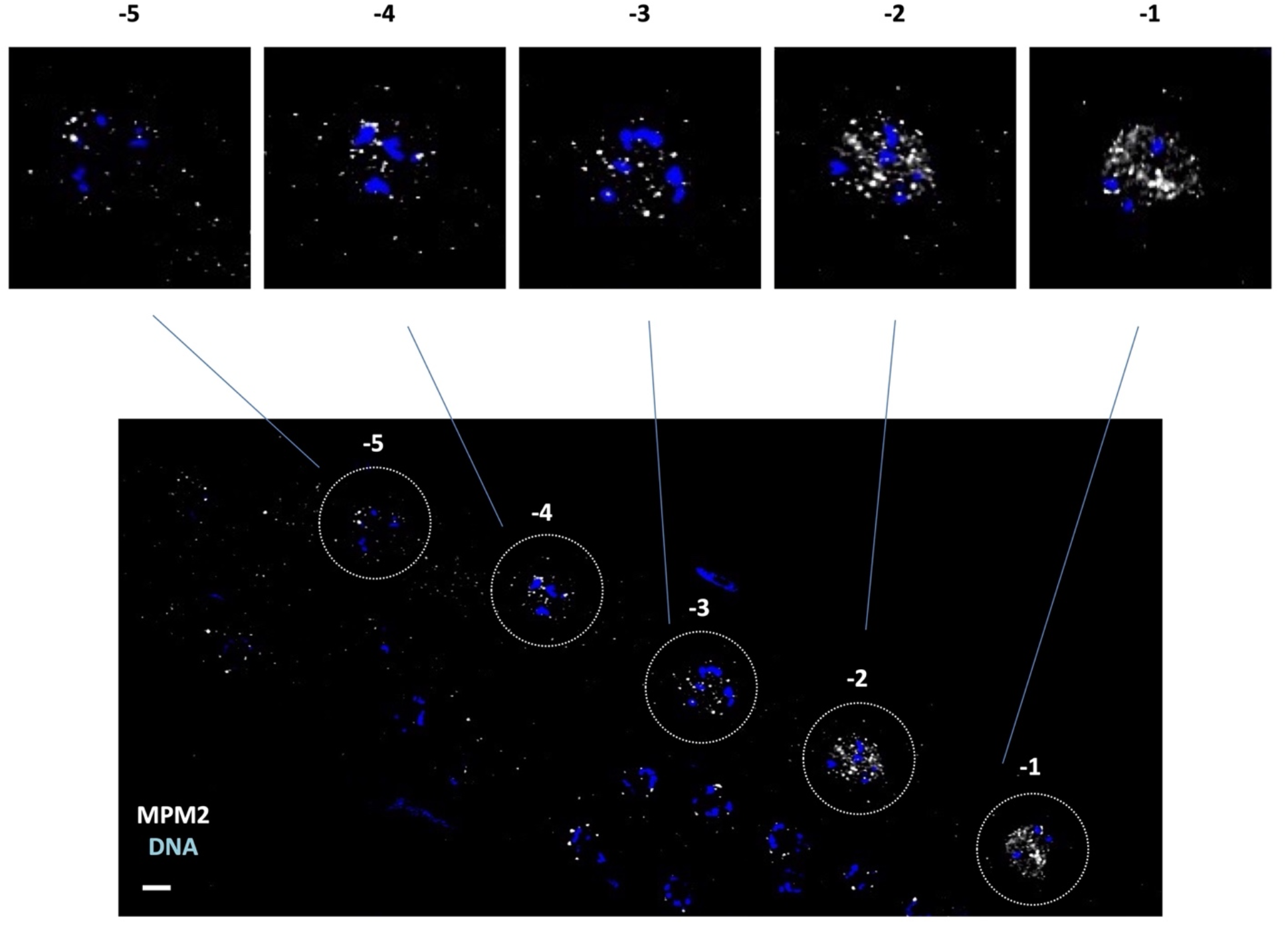
M-phase phosphoproteins gradually increase as oocytes become more proximal. N2 gonads were dissected, fixed, and stained for DNA (blue) and MPM-2 (white). An increase in MPM-2 signal is observed at the proximal oocyte positions in comparison to more distal oocytes. Scale bar represents a length of 2 µm.

**Figure S4:**
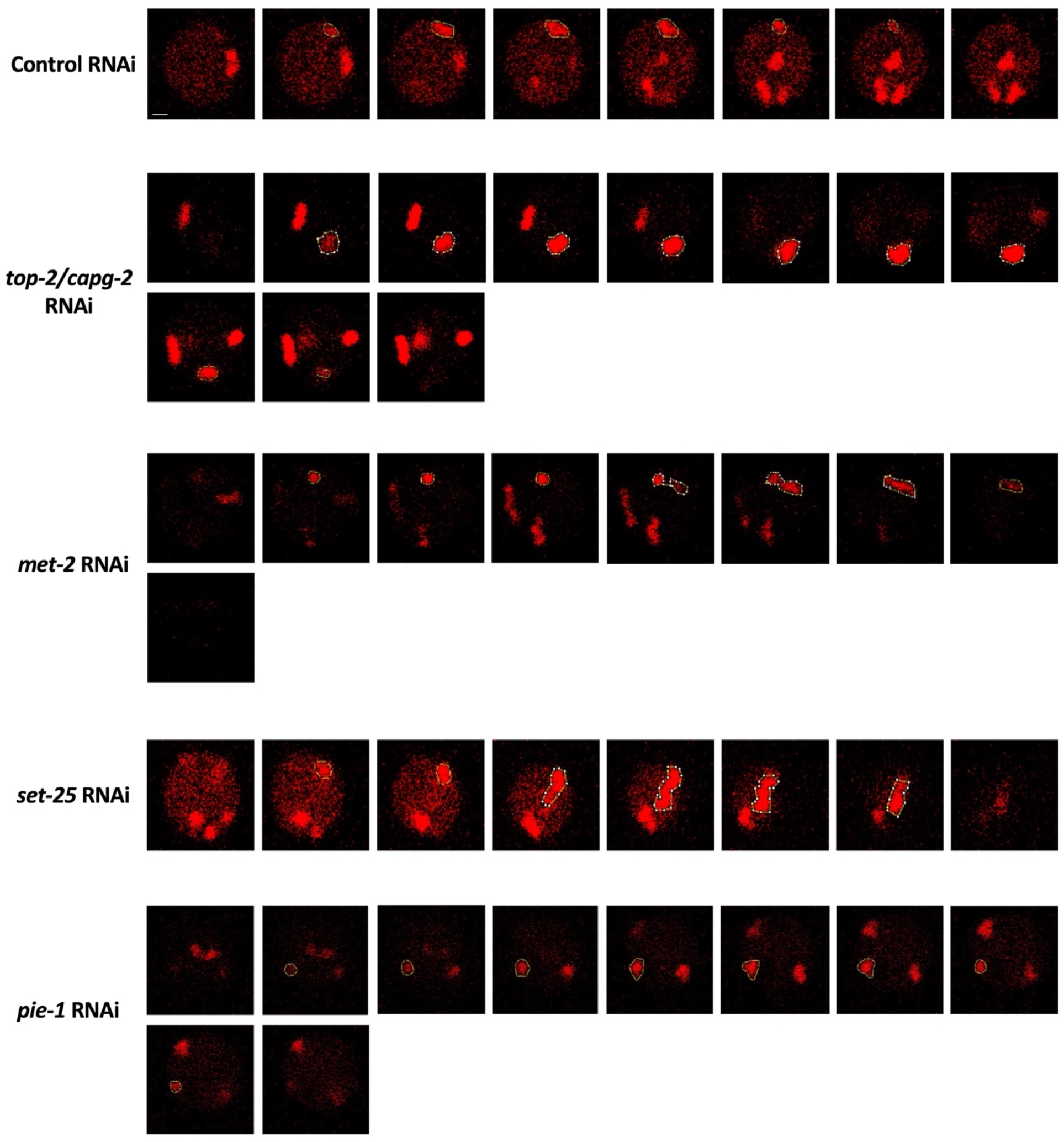
Representative images for bivalent volume measurements. Z-stacks were taken from living oocytes at the −2 position in animals treated with control, *top-2/capg-2*, *met-2*, *set-25*, and *pie-1* RNAi. Shown here are representative images of a bivalent volume measurement from each RNAi treatment. See methods for details on bivalent volume measurement. Scale bar represents a length of 2 µm.

## References

1. Abe S, Nagasaka K, Hirayama Y, Kozuka-Hata H, Oyama M, Aoyagi Y, Obuse C, Hirota T. (2011). The initial phase of chromosome condensation requires Cdk1-mediated phosphorylation of the CAP-D3 subunit of condensin II. Genes Dev. 25(8):863–74.

2. Abe KI, Funaya S, Tsukioka D, Kawamura M, Suzuki Y, Suzuki MG, Schultz RM, Aoki F. (2018). Minor zygotic gene activation is essential for mouse preimplantation development. Proc Natl Acad Sci U S A. 115(29):E6780–E6788.

3. Belew MD, Chien E, Wong M, Michael WM. (2021). A global chromatin compaction pathway that represses germline gene expression during starvation. J Cell Biol. 220(9):e202009197.

4. Bessler JB, Andersen EC, Villeneuve AM. (2010). Differential localization and independent acquisition of the H3K9me2 and H3K9me3 chromatin modifications in the *Caenorhabditis elegans* adult germ line. PLoS Genet. 6(1):e1000830.

5. Bouniol-Baly C, Hamraoui L, Guibert J, Beaujean N, Szöllösi MS, Debey P. (1999). Differential transcriptional activity associated with chromatin configuration in fully grown mouse germinal vesicle oocytes. Biol Reprod. 60(3):580–7.

6. Bowman EA, Bowman CR, Ahn JH, Kelly WG. (2013). Phosphorylation of RNA polymerase II is independent of P-TEFb in the *C. elegans* germline. Development. 140(17):3703–13.

7. Burrows AE, Sceurman BK, Kosinski ME, Richie CT, Sadler PL, Schumacher JM, Golden A. (2006). The *C. elegans* Myt1 ortholog is required for the proper timing of oocyte maturation. Development. 133(4):697–709.

8. Butuči M, Williams AB, Wong MM, Kramer B, Michael WM. (2015). Zygotic Genome Activation Triggers Chromosome Damage and Checkpoint Signaling in *C. elegans* Primordial Germ Cells. Dev Cell. 34(1):85–95.

9. Cassart C, Yague-Sanz C, Bauer F, Ponsard P, Stubbe FX, Migeot V, Wery M, Morillon A, Palladino F, Robert V, Hermand D. (2020). RNA polymerase II CTD S2P is dispensable for embryogenesis but mediates exit from developmental diapause in *C. elegans*. Sci Adv. 6(50):eabc1450.

10. Chan RC, Severson AF, Meyer BJ. (2004). Condensin restructures chromosomes in preparation for meiotic divisions. J Cell Biol. 167(4):613–25.

11. Davis FM, Tsao TY, Fowler SK, Rao PN. (1983). Monoclonal antibodies to mitotic cells. Proc Natl Acad Sci U S A. 80(10):2926–30.

12. De La Fuente R, Viveiros MM, Burns KH, Adashi EY, Matzuk MM, Eppig JJ. (2004). Major chromatin remodeling in the germinal vesicle (GV) of mammalian oocytes is dispensable for global transcriptional silencing but required for centromeric heterochromatin function. Dev Biol. 275(2):447–58.

13. de Morree A, Rando TA. (2023). Regulation of adult stem cell quiescence and its functions in the maintenance of tissue integrity. Nat Rev Mol Cell Biol.

14. Freeman L, Aragon-Alcaide L, Strunnikov A. (2000). The condensin complex governs chromosome condensation and mitotic transmission of rDNA. J Cell Biol. 149(4):811–24.

15. Fujinaga K, Huang F, Peterlin BM. (2023). P-TEFb: The master regulator of transcription elongation. Mol Cell. 83(3):393–403.

16. Gavet O, Pines J. (2010). Progressive activation of CyclinB1-Cdk1 coordinates entry to mitosis. Dev Cell. 18(4):533–43.

17. Ghosh D, Seydoux G. (2008). Inhibition of transcription by the *Caenorhabditis elegans* germline protein PIE-1: genetic evidence for distinct mechanisms targeting initiation and elongation. Genetics.178(1):235–43.

18. Gonzalez I, Molliex A, Navarro P. (2021). Mitotic memories of gene activity. Curr Opin Cell Biol. 69:41–47.

19. Greenstein D. (2005). Control of oocyte meiotic maturation and fertilization. WormBook. 1–12.

20. Guven-Ozkan T, Nishi Y, Robertson SM, Lin R. (2008). Global transcriptional repression in *C. elegans* germline precursors by regulated sequestration of TAF-4. Cell. 135(1):149–60.

21. Hendzel MJ, Wei Y, Mancini MA, Van Hooser A, Ranalli T, Brinkley BR, Bazett-Jones DP, Allis CD. (1997). Mitosis-specific phosphorylation of histone H3 initiates primarily within pericentromeric heterochromatin during G2 and spreads in an ordered fashion coincident with mitotic chromosome condensation. Chromosoma. 106(6):348–60.

22. Hintermair C, Voß K, Forné I, Heidemann M, Flatley A, Kremmer E, Imhof A, Eick D. (2016). Specific threonine-4 phosphorylation and function of RNA polymerase II CTD during M phase progression. Sci Rep. 6:27401.

23. Ito K, Zaret KS. (2022). Maintaining Transcriptional Specificity Through Mitosis. Annu Rev Genomics Hum Genet. 23:53–71.

24. Kim H, Ding YH, Lu S, Zuo MQ, Tan W, Conte D Jr, Dong MQ, Mello CC. (2021). PIE-1 SUMOylation promotes germline fates and piRNA-dependent silencing in *C. elegans*. Elife. 10:e63300.

25. Kimura H, Hayashi-Takanaka Y, Goto Y, Takizawa N, Nozaki N. (2008). The organization of histone H3 modifications as revealed by a panel of specific monoclonal antibodies. Cell Struct Funct. 33(1):61–73.

26. Kobayashi W, Tachibana K. (2021). Awakening of the zygotic genome by pioneer transcription factors. Curr Opin Struct Biol. 71:94–100.

27. Korčeková D, Gombitová A, Raška I, Cmarko D, Lanctôt C. (2012). Nucleologenesis in the *Caenorhabditis elegans* embryo. PLoS One. 7(7):e40290.

28. Maryu G, Yang Q. (2022). Nuclear-cytoplasmic compartmentalization of cyclin B1-Cdk1 promotes robust timing of mitotic events. Cell Rep. 41(13):111870.

29. Mello CC, Schubert C, Draper B, Zhang W, Lobel R, Priess JR. (1996). The PIE-1 protein and germline specification in *C. elegans* embryos. Nature. 382(6593):710–2.

30. Miller, MA, Nguyen, VQ, Lee, MH, Kosinski, M, Schedl, T, Caprioli, RM, Greenstein, D. (2001). A sperm cytoskeletal protein that signals oocyte meiotic maturation and ovulation. Science. 291(5511): 2144–2147.

31. Moore GP, Lintern-Moore S. (1974). A correlation between growth and RNA synthesis in the mouse oocyte. J Reprod Fertil. 39(1):163–6.

32. Moore GP, Lintern-Moore S, Peters H, Faber M. (1974). RNA synthesis in the mouse oocyte. J Cell Biol. 60(2):416–22.

33. Nakamura A, Seydoux G. (2008). Less is more: specification of the germline by transcriptional repression. Development.135(23):3817–27.

34. Navarro-Costa P, McCarthy A, Prudêncio P, Greer C, Guilgur LG, Becker JD, Secombe J, Rangan P, Martinho RG. (2016). Early programming of the oocyte epigenome temporally controls late prophase I transcription and chromatin remodelling. Nat Commun. 7:12331.

35. Palancade B, Bensaude O. (2003). Investigating RNA polymerase II carboxyl-terminal domain (CTD) phosphorylation. Eur J Biochem. 270(19):3859–70.

36. Palozola KC, Donahue G, Liu H, Grant GR, Becker JS, Cote A, Yu H, Raj A, Zaret KS. (2017). Mitotic transcription and waves of gene reactivation during mitotic exit. Science. 358(6359):119–122.

37. Palozola KC, Lerner J, Zaret KS. (2019). A changing paradigm of transcriptional memory propagation through mitosis. Nat Rev Mol Cell Biol. 20(1):55–64.

38. Parsons GG, Spencer CA. (1997). Mitotic repression of RNA polymerase II transcription is accompanied by release of transcription elongation complexes. Mol Cell Biol. 17(10):5791–802.

39. Prescott DM, Bender MA. (1962). Synthesis of RNA and protein during mitosis in mammalian tissue culture cells. Exp Cell Res. 26:260–8.

40. Schaner CE, Deshpande G, Schedl PD, Kelly WG. (2003). A conserved chromatin architecture marks and maintains the restricted germ cell lineage in worms and flies. Dev Cell. 5(5):747–57.

41. Schedl T. (1997). Developmental Genetics of the Germ Line. In: Riddle DL, Blumenthal T, Meyer BJ, Priess JR, editors. C. elegans II. 2nd ed. Cold Spring Harbor (NY): Cold Spring Harbor Laboratory Press; Chapter 10.

42. Schisa JA, Pitt JN, Priess JR. (2001). Analysis of RNA associated with P granules in germ cells of *C. elegans* adults. Development. 128(8):1287–98.

43. Schultz RM, Stein P, Svoboda P. (2018). The oocyte-to-embryo transition in mouse: past, present, and future. Biol Reprod. 99(1):160–174.

44. Seydoux G, Dunn MA. (1997). Transcriptionally repressed germ cells lack a subpopulation of phosphorylated RNA polymerase II in early embryos of *Caenorhabditis elegans* and Drosophila melanogaster. Development. 124(11):2191–201.

45. Seydoux G, Mello CC, Pettitt J, Wood WB, Priess JR, Fire A. (1996). Repression of gene expression in the embryonic germ lineage of *C. elegans*. Nature. 382(6593):713–6.

46. Smith R, Susor A, Ming H, Tait J, Conti M, Jiang Z, Lin CJ. (2022). The H3.3 chaperone Hira complex orchestrates oocyte developmental competence. Development. 149(5):dev200044.

47. Swygert SG, Tsukiyama T. (2019). Unraveling quiescence-specific repressive chromatin domains. Curr Genet. 65(5):1145–1151.

48. Taylor JH. (1960). Nucleic acid synthesis in relation to the cell division cycle. Ann N Y Acad Sci. 90:409–21.

49. Tora L, Vincent SD. (2021). What defines the maternal transcriptome? Biochem Soc Trans. 49(5):2051–2062.

50. Towbin BD, González-Aguilera C, Sack R, Gaidatzis D, Kalck V, Meister P, Askjaer P, Gasser SM. (2012). Step-wise methylation of histone H3K9 positions heterochromatin at the nuclear periphery. Cell. 150(5):934–47.

51. Walker AK, Boag PR, Blackwell TK. (2007). Transcription reactivation steps stimulated by oocyte maturation in *C. elegans*. Dev Biol. 304(1):382–93.

52. Wang JT, Seydoux G. (2013). Germ cell specification. Adv Exp Med Biol. 757:17–39.

53. Wong MM, Belew MD, Kwieraga A, Nhan JD, Michael WM. (2018). Programmed DNA Breaks Activate the Germline Genome in *Caenorhabditis elegans*. Dev Cell. 46(3):302–315.

54. Zhang F, Barboric M, Blackwell TK, Peterlin BM. (2003). A model of repression: CTD analogs and PIE-1 inhibit transcriptional elongation by P-TEFb. Genes Dev. 17(6):748–58.

55. Zuccotti M, Piccinelli A, Giorgi Rossi P, Garagna S, Redi CA. (1995). Chromatin organization during mouse oocyte growth. Mol Reprod Dev. 41(4):479–85.

